# Modeling human limb skeletal development using human pluripotent stem cell-derived skeletal assembloids

**DOI:** 10.1101/2025.01.09.631479

**Authors:** Tomoka Takao, Tatsunori Osone, Kohei Sato, Daisuke Yamada, Yuki Fujisawa, Masaya Hagiwara, Eiji Nakata, Toshifumi Ozaki, Junya Toguchida, Takeshi Takarada

## Abstract

Despite recent advances in pluripotent stem cell-based approaches to induce skeletal cells, recapitulating human limb skeletal development in terms of structure and longitudinally oriented growth remains an unresolved challenge. Here, we report a method to differentiate human pluripotent stem cells into region-specific skeletal organoids harboring GDF5^+^PRG4^+^ interzone/articular chondrocyte progenitors (IZ/ACPs) and SP7^+^ growth plate chondrocytes (GPCs) via PRRX1^+^ limb-bud mesenchymal cells. Comparative analysis demonstrated marked similarities of IZ/ACP and GPC organoids to the human embryonic limb, and graft fate and regenerative capacity *in vivo* were further characterized. We also mimicked the limb skeletal developmental process in a spatially structured manner by vertically positioning two IZ/ACP organoids at both ends of a GPC organoid to generate a human skeletal assembloid. Notably, this human skeletal assembloid recapitulated endochondral ossification with longitudinal skeletal growth upon transplantation. In summary, our study provides a novel research platform for human limb skeletal development and disease.

## INTRODUCTION

Understanding human skeletal development is essential not only to advance bone physiology, e.g., providing a weight-bearing framework, regulating endocrine functions, and supporting hematopoiesis ^1^, but also to address clinical challenges, including osteoarthritis ^2,3^, nonunion fractures ^4^ in the elderly, and congenital skeletal disorders ^5^, which remain unresolved by conventional therapies. Although ethical constraints on research into human skeletal development have long existed, advancements in human pluripotent stem cells (hPSCs) have transformed this field. With their self-renewal and pluripotency, hPSCs provide experimental models that mimic skeletal development by navigating the brachiating landscape and efficiently differentiating intermediate stem/progenitor cells into desired cell types for regenerative medicine and recapitulation of congenital skeletal disorders ^6^. In mammals, the skeleton is derived from three distinct embryonic sources according to the anatomical region. The axial skeleton (e.g., vertebrae and ribs) is specified from the paraxial mesoderm, the cranial skeleton (e.g., facial and cranial bone) is derived from neural crest, whereas the appendicular skeleton (e.g., limbs) arises from the lateral plate mesoderm (LPM) ^7^. Moreover, ossification during skeletal development is mediated by two distinct processes: intramembranous ossification and endochondral ossification. In intramembranous ossification, mesenchymal cells directly differentiate into osteoblasts that are responsible for bone formation ^8^. Conversely, in endochondral ossification, a cartilaginous template (anlagen) is initially established, which is subsequently replaced by bone ^9^. Recent studies employed hPSCs to recapitulate skeletal lineages, successfully inducing skeletal cells or organoids (three-dimensional (3D) structures grown from stem cells that mimic certain aspects of organs) that participate in intramembranous ^10–14^ or endochondral ^15–21^ ossification. However, replicating processes that emulate skeletal morphology with skeletal growth using hPSCs remains an unresolved challenge.

Development of limb skeletal structures is initiated by condensation of PRRX1^+^ limb-bud mesenchymal (LBM) cells, followed by formation of skeletal anlagen composed of immature chondrocytes ^22^. The joint interzone develops at the position of the presumptive joint and precedes articular cartilage differentiation and joint cavitation. Immature chondrocytes outside the interzone undergo hypertrophy at the mid-diaphysis as the first step of endochondral ossification ^22^. Vasculature invades hypertrophic chondrocytes and subsequently forms the bone marrow cavity and growth plate chondrocytes (GPCs)/cartilage, which support longitudinal skeletal growth of limbs until puberty ^9,22^. To replicate these complex human limb developmental processes, induction of LBM cells from hPSCs is essential. Recently, we successfully generated PRRX1^+^ LBM cells from hPSCs in a stable and scalable manner by employing PRRX1-tdTomato reporter hPSCs ^23,24^. This achievement laid the foundation for induction of intermediate structures, such as the interzone and growth plate cartilage, from hPSCs. In this study, we developed a protocol to generate distinct intermediate organoids, characterized by interzone/articular chondrocyte progenitors (IZ/ACPs) or GPCs, from ontogenetically-defined PRRX1^+^ LBM cells derived from hPSCs via establishing lineage-specific tdTomato reporter hPSC lines. Additionally, by employing a 3D assembly strategy for these intermediate tissue constructs, we successfully recapitulated spatial developmental processes, thereby reproducing skeletal growth along the longitudinal axis.

## RESULTS

### Generation of ontogenetically-defined IZ/ACP and GPC organoids from hPSCs

In our previous study, we induced and expanded PRRX1^+^ limb-bud-like mesenchymal cells (expandable limb-bud-like mesenchymal cells, ExpLBM cells) from hPSCs ^23^. This protocol begins with differentiation into the mid-primitive streak and LPM, followed by induction of LBM cells, which have chondrogenic capacity in the presence of BMP4, TGFβ1, and GDF5 (Figure S1A). The expression level of each marker gene (pluripotency, LPM, LBM, and chondrocyte markers) clearly revealed the sequential induction of genes representative of each differentiation stage (Figure S1B). Chromatin accessibility, which is epigenetically regulated by histone modifications such as acetylation and methylation, determines the access of transcription factors and underlies cell type-specific gene expression ^25,26^. Therefore, we investigated cell specificity during the stepwise induction protocol by analyzing chromatin accessibility using ATAC-seq and histone modifications using CUT&Tag. *NANOG, HAND1*, and *PRRX1* are specific markers of pluripotent, LPM, and LBM cells, respectively, and mRNA expression of these genes was specific to each developmental stage. When the 5’ flanking regions of these genes were analyzed using ATAC-seq and CUT&Tag with antibodies against H3K27Ac and H3K27me3, peaks of ATAC-seq and H3K27Ac were observed at time points corresponding to the mRNA expression peaks. By contrast, H3K27me3 peaks were inversely correlated with these patterns (Figure S1C). Furthermore, we assessed whether our ontogenetically-defined induction protocol for ExpLBM cells recapitulates formation of the forelimb or hindlimb. Reanalysis of single-cell RNA-sequencing (scRNA-seq) data of human forelimb and hindlimb buds ^27^ revealed that *TBX5* was highly expressed in forelimb buds, while *PITX1* and *ISL1* were enriched in hindlimb buds, consistent with previous mouse studies ^28^ (Figure S1D). *PITX1* and *ISL1*, but not *TBX5*, were expressed during stepwise induction processes from hPSCs to ExpLBM cells (Figure S1D). Under these conditions, high peak patterns of ATAC-seq and H3K27Ac were found in *ISL1* and *PITX1*, but not in *TBX5*. On the other hand, H3K27me3 peaks were found in *TBX5*, but not in *ISL1* and *PITX1* (Figure S1E). These results indicate that our induction protocols generate ontogenetically-defined LBM cells that recapitulate hindlimb bud initiation.

Limb skeletal development begins with condensation of LBM cells, forming a cartilage template. This template subsequently differentiates into two distinct lineages: articular permanent cartilage, which develops through intermediate GDF5-expressing interzone cells, and growth plate cartilage comprising resting, proliferative, and SP7-expressing hypertrophic chondrocytes, which drives longitudinal bone growth. Our previous studies generated cartilage tissues from ExpLBM cells by 30 ng/mL BMP4 and 10 ng/mL TGFβ ^23,24^. However, RNA-seq analysis of cartilage generated using this protocol revealed expression of markers for both interzone/articular cartilage and growth plate cartilage (Figure S1B). In addition, subcutaneous transplantation of this cartilage into NOD-SCID mice revealed that some transplanted cartilage maintained hyaline cartilaginous-like tissues, but some showed ossification with columnar chondrocytes and spongy bone at 8 weeks (Figure S1F). Thus, our previous chondrogenic induction protocol from ExpLBM cells does not seem to selectively induce each lineage for interzone/articular cartilage or growth plate cartilage. To develop a lineage-selective induction protocol for ExpLBM cells into interzone/articular cartilage or growth plate cartilage, we established GDF5-tdTomato reporter human induced pluripotent stem cells (GDF5^3’tdTomato^ hiPSCs) and SP7-tdTomato reporter human embryonic stem cells (SP7^3’tdTomato^ hESCs) to visualize GDF5 (interzone marker)^+^ or SP7 (hypertrophic chondrocyte marker)^+^ cells. hPSCs were electroporated with the targeting vector harboring a homology arm and IRES-tdTomato-PGK-Neo cassette, and the PX459-gRNA genome-editing vector. Electroporated cells were selected with puromycin and G418, followed by cell cloning to isolate 3’tdTomato reporter hPSCs. Single colonies were expanded, and genomic DNA was extracted to identify correct insertion of the knock-in reporter cassette. PCR results showed that the selected clone harbored a homozygous knock-in reporter cassette (Figure S2A). These reporter clones of GDF5^3’tdTomato^ hiPSCs or SP7^3’tdTomato^ hESCs were chosen for induction experiments.

Next, ExpLBM cells derived from GDF5^3’tdTomato^ hiPSCs or SP7^3’tdTomato^ hESCs were condensed into aggregates (step 1) and subsequently differentiated into chondrocytes using BMP4, TGFβ1, and FGF2 (step 2). This was followed by treatment with 18 combinations of chemicals in the presence of 30 ng/mL BMP4, 10 ng/mL TGFβ1, and 10 ng/mL GDF5 for 14 days (Figure 1A). Among the combinations tested, addition of PD0325901 (a MEK inhibitor) in the presence of BMP4 and TGFβ1 was necessary to induce GDF5-tdTomato expression. By contrast, high concentrations of BMP4 and TGFβ1 (300 and 100 ng/mL, respectively) effectively induced SP7-tdTomato expression (Figure 1B and S2B). Principal component analysis revealed distinct differential gene expression profiles between pluripotent cells (day 0), LPM cells (day 2), LBM cells (day 4), ExpLBM cells, and cell aggregates treated with 30 ng/mL BMP4 and 10 ng/mL TGFβ1 (conventional); 30 ng/mL BMP4, 10 ng/mL TGFβ1, and PD0325901 (hereafter referred to as the IZ/ACP organoid strategy); or 300 ng/mL BMP4 and 100 ng/mL TGFβ1 (hereafter referred to as the GPC organoid strategy) (Figure 1C). Notably, RNA-seq analysis clearly demonstrated that interzone/articular cartilage markers (e.g., *GDF5, PRG4,* and *BARX1*) were enriched in IZ/ACP organoids, whereas growth plate cartilage markers (e.g., *RUNX2, SP7, IHH,* and *COL10A1*) were highly expressed in GPC organoids (Figure 1D). We next assessed the expression pattern of each marker by immunohistochemistry in ExpLBM cell aggregates and IZ/ACP and GPC organoids. While SOX9 and aggrecan (ACAN) were consistently expressed in all tested aggregates, PRG4 and type I collagen (COL1) were detected exclusively in IZ/ACP organoids. By contrast, RUNX2, type X collagen (COL10), and SP7 were observed only in GPC organoids (Figure 1E). We also confirmed that two different types of hPSCs (802-3G hiPSCs and SEES4 hESCs) could established similar expression profiles by performing real-time quantitative RT-PCR (RT-qPCR) analysis (Figure S2C).

**Figure 1:**
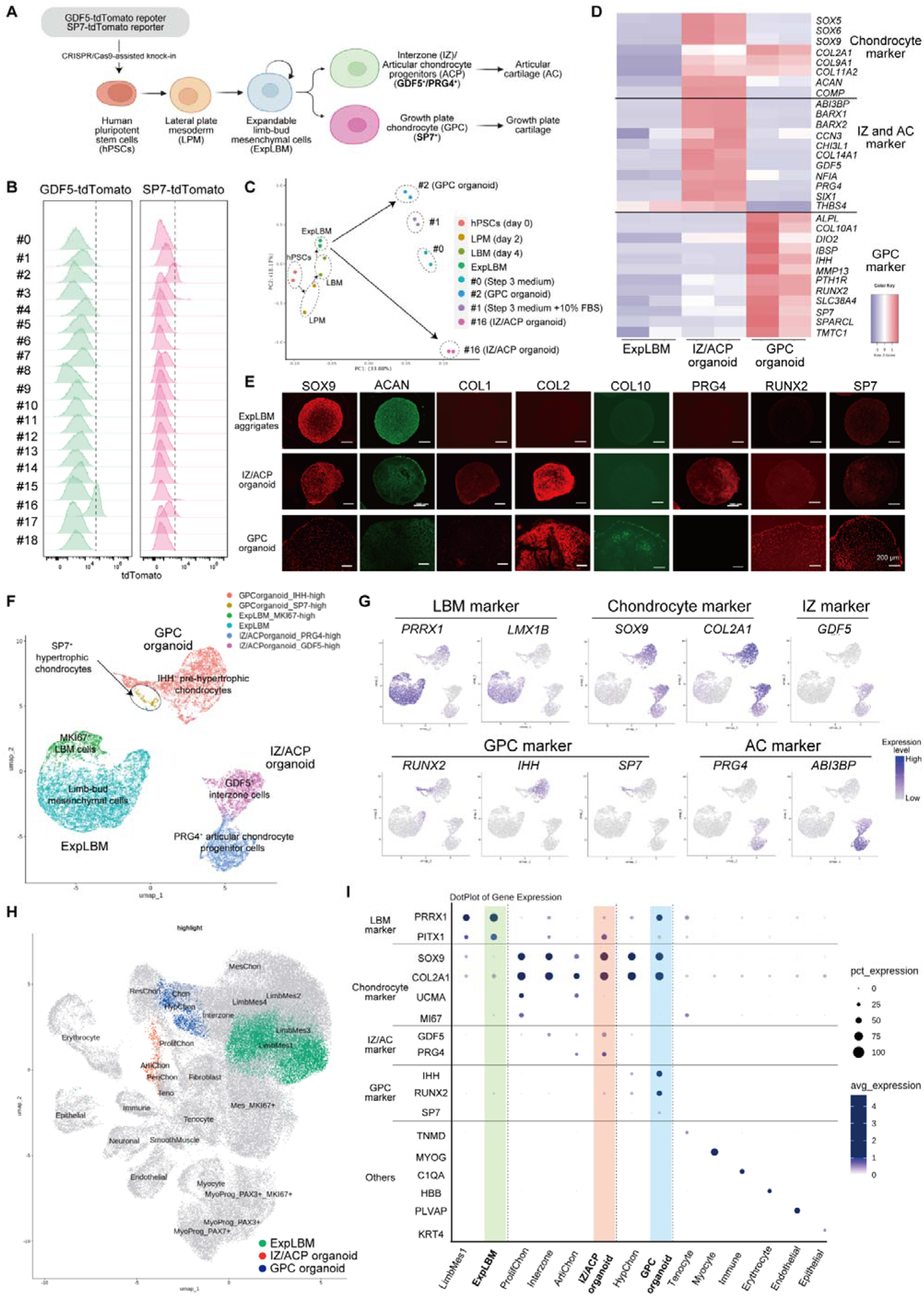
*In vitro* stepwise induction of hPSC-derived LBM cells into IZ/ACP and GPC organoids. (A) Overview of the stepwise induction and differentiation of human LBM cells, followed by chondrocyte lineages, which closely mimics embryonic development. The optimal conditions were determined for ExpLBM cells to induce IZ/ACP and GPC organoids using GDF5- and SP7-tdTomato reporters, respectively. (B) tdTomato fluorescence intensity in GDF5^3’tdTomato^ or SP7^3’tdTomato^ reporter-derived ExpLBM cells treated with 18 combinations of chemicals and cytokines for 14 days in the presence of 30 ng/mL BMP4 and 10 ng/mL TGFβ1. #0, none; #1, 10% FBS; #2, 300 ng/mL BMP4, 100 ng/mL TGFβ1, and 10% FBS; #3, CHIR99021 (GSK3β inhibitor and WNT activator); #4, C59 (PORCN inhibitor and WNT inhibitor); #5, XAV-939 (tankyrase 1 inhibitor and WNT inhibitor); #6, LDN-193189 (ALK2/3 inhibitor); #7, A 83-01 (ALK4/5/7 inhibitor); #8, SAG 21k (hedgehog signaling activator); #9, Vismodegib (hedgehog signaling inhibitor); #10, MHY1485 (mTOR activator); #11, VO-OHpic (PTEN inhibitor and PI3K/mTOR activator); #12, SF1670 (PTEN inhibitor and PI3K/mTOR activator); #13, rapamycin (mTOR inhibitor); #14, ATRA (RA signaling activator); #15, MK-206 (AKT inhibitor); #16, PD0325901 (MEK inhibitor); #17, PIK-90 (PI3K inhibitor); and #18, DAPT (γ-secretase inhibitor and Notch inhibitor). (C–D) Bulk RNA-seq transcriptome analysis of hPSCs (day 0), LPM cells (day 2), LBM cells (day 4), ExpLBM cells (passage numbers 4–5), and aggregates treated under the following conditions: (i) 30 ng/mL BMP4 and 10 ng/mL TGFβ1 (condition #0), (ii) 30 ng/mL BMP4, 10 ng/mL TGFβ1, and 10% FBS (condition #1), (iii) 30 ng/mL BMP4, 10 ng/mL TGFβ1, and 500 nM PD-0325901 (condition #16, referred to as IZ/ACP organoids), and (iv) 300 ng/mL BMP4, 100 ng/mL TGFβ1, and 10% FBS (condition #2, referred to as GPC organoids). n = 2 biologically independent experiments. Principal component analysis (C) and a heatmap of the gene expression level of each marker (D). (E) Immunohistological analysis of ExpLBM aggregates, IZ/ACP organoids, and GPC organoids with antibodies against SOX9, ACAN, COL1, COL2, COL10, PRG4, RUNX2, and SP7. (F) UMAP embedding of scRNA-seq data from ExpLBM cells (9741 cells), IZ/ACP organoids (5090 cells), and GPC organoids (4660 cells). UMAP clustering identified six clusters, including LBM cells (non-dividing) and MKI67^+^ LBM cells (dividing) in ExpLBM cells, GDF5^+^ interzone cells and PRG4^+^ articular chondrocyte progenitors in IZ/ACP organoids, and IHH^+^ pre-hypertrophic chondrocytes and SP7^+^ hypertrophic chondrocytes in GPC organoids. (G) Feature plot of each marker gene in UMAP of integrated scRNA-seq datasets from ExpLBM cells, IZ/ACP organoids, and GPC organoids. (H) UMAP of integrated scRNA-seq profiles of ExpLBM cells, IZ/ACP organoids, and GPC organoids with human embryonic limb cells at PCW 5–9 (119,870 cells). Cells are colored by origin: human embryonic limb cells (gray), ExpLBM cells (green), IZ/ACP organoids (red), and GPC organoids (blue). The original datasets of human embryonic limb cells are from Zhang et al. (I) Dot plots showing expression of cell type-specific genes in each cluster of scRNA-seq analysis of human embryonic limb cells and scRNA-seq profiles of ExpLBM cells, IZ/ACP organoids, and GPC organoids.

To further analyze lineage specification and its heterogeneity, we performed scRNA-seq of 9741 cells from ExpLBM cell aggregates, 5090 cells from IZ/ACP organoids, and 4660 cells from GPC organoids. Uniform manifold approximation and projection (UMAP) dimensionality identified two clusters in ExpLBM cells [LBM cells (non-dividing) and MKI67^+^ LBM cells (dividing)], two clusters in IZ/ACP organoids (GDF5^+^ interzone cells and PRG4^+^ articular chondrocyte progenitors), and two clusters in GPC organoids (IHH^+^ pre-hypertrophic chondrocytes and SP7^+^ hypertrophic chondrocytes) based on gene expression patterns (Figures 1F and 1G,). To evaluate the similarity between these hPSC-derived cells and cells from human embryos, we integrated their scRNA-seq expression profiles with data from the human limb developmental process ^27^. Our analysis revealed that ExpLBM cells, IZ/ACP organoids, and GPC organoids most closely resembled limb-bud mesenchyme (LimbMes1), articular chondrocytes, and hypertrophic chondrocytes observed during human limb development, respectively (Figure 1H). Dot plots from scRNA-seq analyses of ExpLBM cells, IZ/ACP organoids, and GPC organoids confirmed their identities as limb-bud mesenchyme, interzone and articular chondrocytes, and hypertrophic chondrocytes, respectively, by comparing the expression of cell type-specific genes in each cluster with those observed in scRNA-seq analyses of human embryonic limb cells (Figure 1I). Furthermore, KEGG pathway enrichment analysis was performed of genes exhibiting significant differential expression identified through pseudotime trajectory analysis using scRNA-seq datasets of human embryonic limbs, with LimbMes1 designated the initial state. This analysis revealed that, unlike differentiation into GPCs, the developmental transition from LBM cells to articular chondrocytes was characterized by suppression of MAPK signaling (Figures S3A, S3B, S3C and S3D). These findings indicate that our differentiation strategy from hPSCs into two distinct chondrocyte lineages, including IZ/ACP and GPC organoids, recapitulates the human skeletal developmental process of the hindlimb bud.

### Assessment of the graft fate and regenerative capacity of IZ/ACP and GPC organoids

Next, we investigated graft fate by subcutaneously transplanting these organoids into NOD-SCID mice. Transplanted IZ/ACP organoids remained Safranin O staining^+^ hyaline cartilaginous-like tissues without ossification up to 3 months post-transplantation, whereas transplanted GPC organoids exhibited ossification, with columnar chondrocytes and spongy bone formation observed at 3 months (Figure 2A). Furthermore, cells of mouse articular cartilage exhibit smooth surfaces, as observed via scanning electron microscopy (SEM), and possess simple intracellular organelle structures, as observed via transmission electron microscopy (TEM). This phenotype closely resembled that of transplanted IZ/ACP organoid-derived cartilaginous tissues. By contrast, cells of mouse growth plate cartilage are characterized by an abundance of cellular projections (observed via SEM) and are rich in rough endoplasmic reticulum and Golgi apparatus (observed via TEM). This appearance was strikingly similar to that of transplanted GPC organoid-derived cartilaginous tissues (Figure 2B). Indeed, immunohistochemical analysis revealed that transplanted IZ/ACP organoid-derived cartilaginous tissues were positively stained for type II collagen (COL2) and ACAN but did not express pre-/hypertrophic chondrocyte markers, such as RUNX2 and SP7, or the endothelial cell marker endomucin (EMCN). By contrast, transplanted GPC organoid-derived cartilaginous tissues expressed not only COL2 and ACAN but also RUNX2 and SP7, and contained EMCN^+^ cells (Figure 2C).

**Figure 2:**
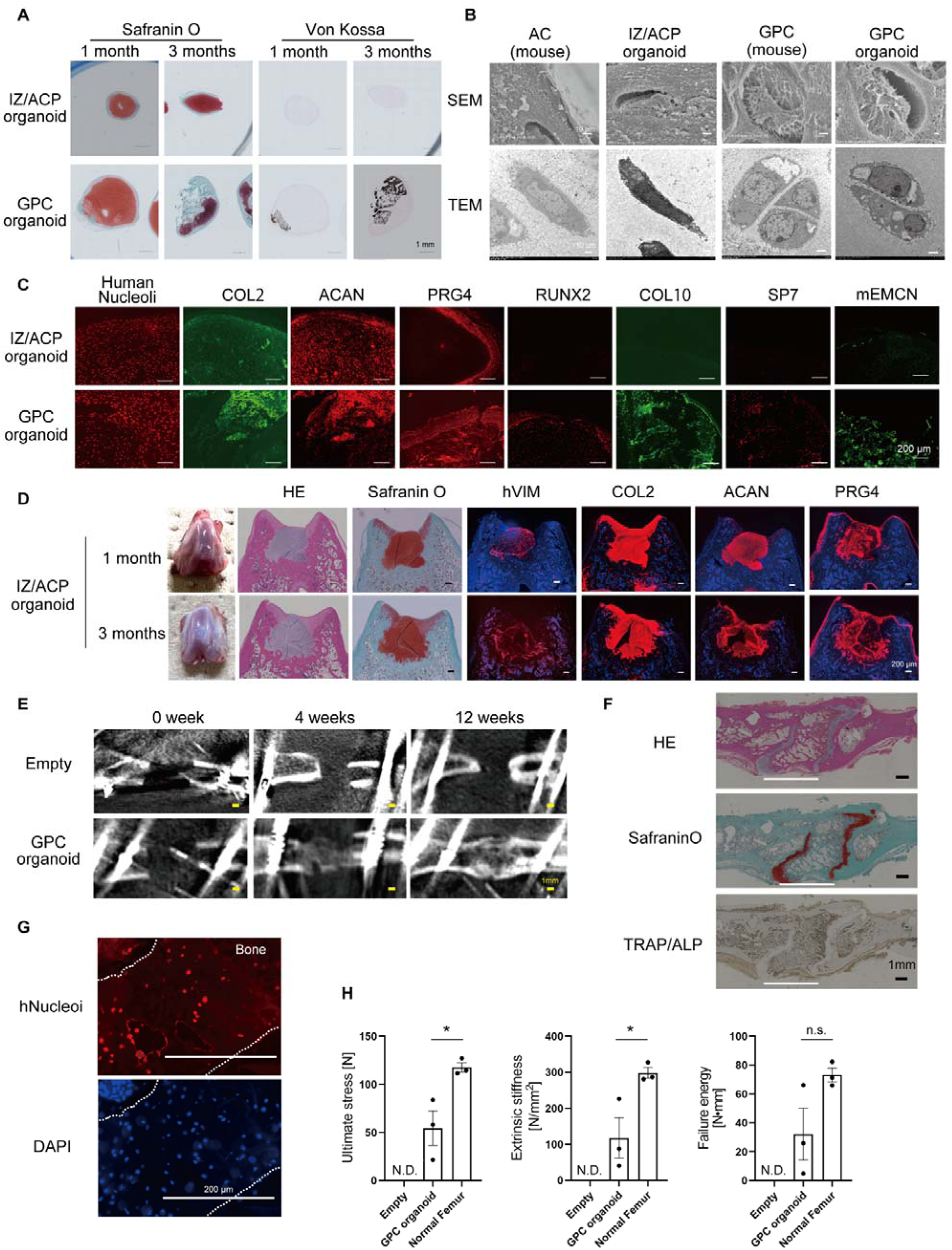
Assessment of the graft fate and regenerative capacity of IZ/ACP and GPC organoids. (A–C) Subcutaneous transplantation of IZ/ACP or GPC organoids into NOD-SCID mice. Their engraftment was histologically assessed at 12 weeks post-transplantation by Safranin O and Von Kossa staining (A), SEM and TEM (B), and immunohistological analysis (C). (D) Transplantation of ExpLBM-derived IZ/ACP organoids into osteochondral defects of articular cartilage in SCID rats. The knee joints were harvested at 1 and 3 months post-transplantation. Histological examination was performed of osteochondral defects engrafted with IZ/ACP organoids. At 1 and 3 months post-transplantation, knee joints were fixed, and tissue sections were stained with HE or Safranin O or an antibody against hVIM, COL2, ACAN, or PRG4. Representative images are displayed. (E) Engraftment of GPC organoids into critical-sized femoral defects of SCID rats was assessed at 4 and 12 weeks post-transplantation using µCT analysis. (F–G) Engraftment of GPC organoids into critical-sized femoral defects of SCID rats was histologically assessed at 8 weeks post-transplantation using HE, Safranin O, and ALP/TRAP staining (F), as well as immunostaining with an anti-human nucleoli antibody (G). (H) Bone strength in critical-sized femoral defects was assessed at 12 weeks post-transplantation using a three-point bending test. *p < 0.05, **p < 0.01, ***p < 0.001, ****p < 0.0001. N.D., not detected.

To demonstrate the regenerative efficacy of IZ/ACP organoids *in vivo*, they were transplanted into osteochondral defects created in knee joint articular cartilage of X-linked severe combined immunodeficiency (SCID) rats. At 1 and 3 months post-transplantation, IZ/ACP organoids were successfully engrafted into osteochondral defects, as evidenced by human vimentin (hVIM) expression. These organoids produced extracellular matrix that was positive for Safranin O staining and immunohistochemical staining for COL2 and ACAN. Notably, PRG4, a marker of the superficial zone of articular cartilage, was specifically expressed in the joint-side surface of the graft (Figure 2D). During fracture healing, the cartilaginous callus that forms at the injury site undergoes endochondral ossification, facilitating bone repair ^29^. To evaluate the regenerative capacity of GPC organoids, we transplanted them into critical-sized femur defects of SCID rats. Periodic observation of mineralized bone using micro-computed tomography (µCT) revealed that, in the absence of transplantation, bone union did not occur, resulting in a condition resembling nonunion. By contrast, transplantation of GPC organoids led to bone formation at the defect site, which was observed by day 28 post-transplantation, with complete bone union achieved by day 56 (Figure 2E). Histological analysis further confirmed active bone formation at the transplantation site, with remnants of Safranin O-stained hyaline cartilaginous-like matrix (Figure 2F). Human-specific nucleoli staining demonstrated that newly formed bone was predominantly graft-derived (Figure 2G). Finally, bone strength at the fracture site was assessed 3 months post-transplantation using a three-point bending test. Although the values did not reach those of intact femurs without defects, significant recovery of ultimate stress (N), extrinsic stiffness (N/mm²) and failure energy (N mm) was observed in GPC organoid-transplanted femurs (Figure 2H). Collectively, these results demonstrate that IZ/ACP organoids exhibit characteristics of IZ/ACPs, which generate permanent articular cartilage, whereas GPC organoids display characteristics of GPCs, driving endochondral ossification.

### Generation of skeletal assembloids with IZ/ACP and GPC organoids to recapitulate human limb skeletal development

The process of skeletal formation begins with condensation of LBM cells, leading to formation of cartilage anlagen. Subsequently, this structure develops into the cartilage containing growth plate chondrocytes, contributing to elongation of bone along its longitudinal axis through endochondral ossification, which is flanked by the interzone, the region that eventually gives rise to joints (Figure 3A) ^22^. The generation of two types of organoids with characteristics of IZ/ACPs or GPCs enabled reconstruction of this skeletal complex *in vitro*. To this end, we fused IZ/ACP and GPC organoids to produce a skeletal assembloid. To facilitate visualization, we inserted a ubiquitous GFP or Kusabira Orange (KuO) marker into hiPSCs using a piggyBac transposon system, followed by generation of IZ/ACP or GPC organoids, respectively (Figure 3B). Two GFP-labeled IZ/ACP organoids were positioned vertically at both ends of a KuO-labeled GPC organoid within a low-melting point agarose gel. By day 7, the three organoids had fused within the wells (Figure 3C), forming a skeletal complex, which was subsequently subcutaneously transplanted into NOD-SCID mice. At 1 and 6 months post-transplantation, excised tissues retained expression of GFP and KuO (Figure 3D). Notably, significant longitudinal growth of the skeletal assembloid was observed over time post-transplantation (Figure 3E). Furthermore, μCT analysis at 6 months post-transplantation revealed ossification in the central region (Figure 3F). Histological analysis of transplanted skeletal assembloids at 1 month post-transplantation demonstrated that GFP^+^ regions were observed at the peripheral edges of the tissue, while KuO^+^ regions were localized in the central area. ACAN was detected throughout the entire tissue, whereas PRG4 was predominantly observed in the peripheral regions. COL10 and RUNX2 were detected in the KuO^+^ central region of the tissue. By contrast, expression of CTSK or EMCN was not observed (Figure 3G). At 6 months post-transplantation, GFP^+^ regions remained localized at the peripheral edges of the transplanted tissue, as observed at 1 month post-transplantation. These regions were positive for Safranin O staining, ACAN, and PRG4. Meanwhile, in KuO^+^ regions at 6 months post-transplantation, a columnar structure characteristic of growth plate cartilage positive for COL10 and RUNX2 was observed. Additionally, bone marrow-like structures were identified in the central region of the tissue construct, along with CTSK^+^ cells and vascular-like structures expressing EMCN (Figure 3H). Overall, our results demonstrate that the skeletal assembloid, which comprises IZ/ACP and GPC organoids, recapitulates longitudinal bone growth upon transplantation due to induction of primary endochondral ossification in its center.

**Figure 3:**
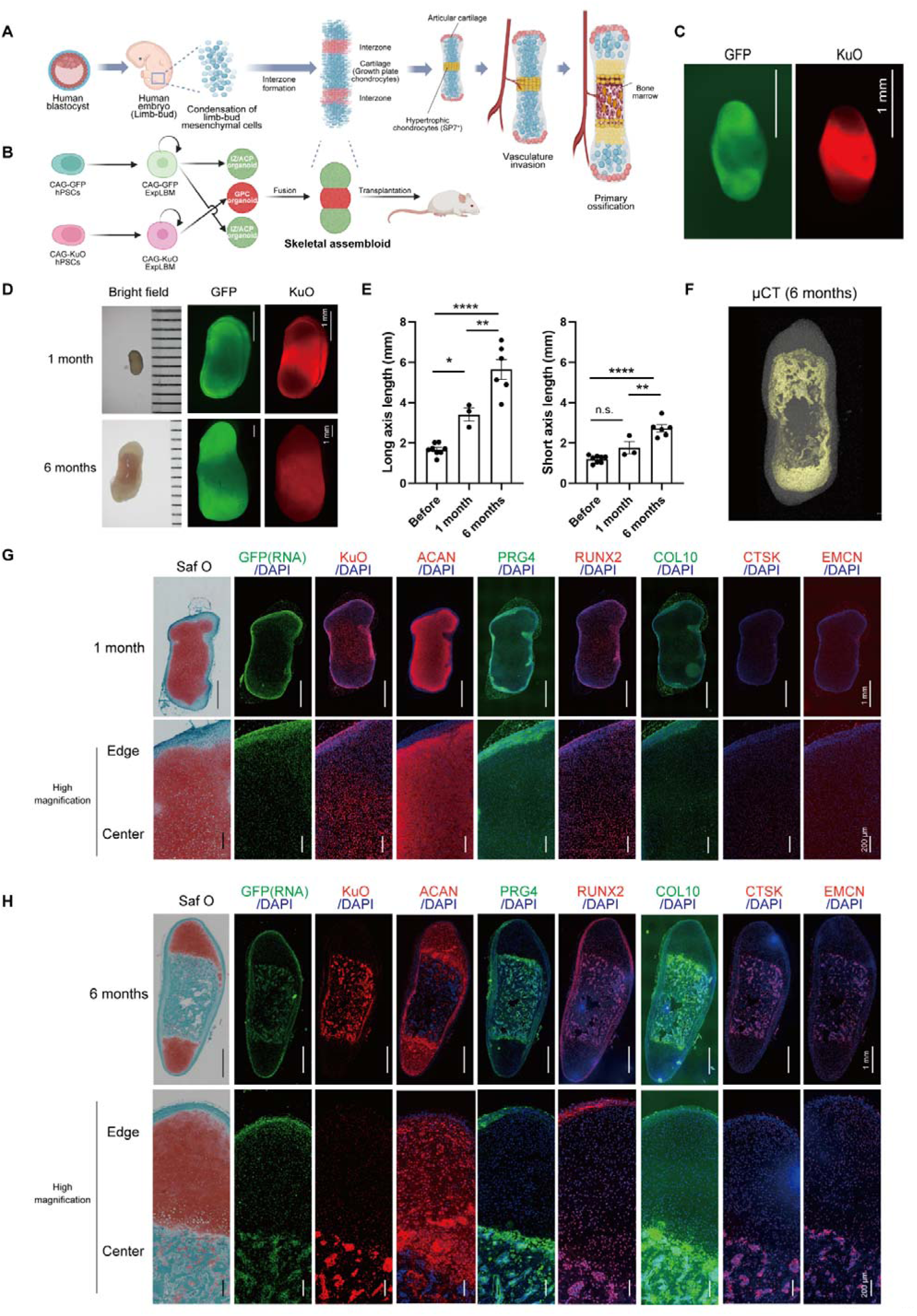
Generation of skeletal assembloids from IZ/ACP and GPC organoids to recapitulate human limb skeletal development. (A) Schematic drawing of skeletal development in the limb. Skeletogenesis begins with LBM cell condensation, followed by formation of skeletal anlagen composed of immature chondrocytes. The joint interzone develops at the position of the presumptive joint and precedes articular cartilage differentiation and joint cavitation. Immature chondrocytes outside the interzone undergo hypertrophy at the mid-diaphysis as the first step of endochondral ossification. Vasculature invades hypertrophic chondrocytes and subsequently forms the bone marrow cavity and growth plate cartilage, which support growth and elongation of the limbs until puberty, when their closure halts any further skeletal growth. (B) Graphical illustration of the approach to generate assembloids from GFP-labeled IZ/ACP organoids and KuO-labeled GPC organoids. (C) Fluorescent images of a typical example of a skeletal assembloid fabricated with a KuO-labeled GPC organoid sandwiched by two GFP-labeled IZ/ACP organoids, followed by culture for 7 days. (D) Subcutaneous transplantation of a skeletal assembloid into NOD-SCID mice. Representative bright-field and fluorescent images of the assembloid at 1 and 6 months post-transplantation. (E) Long and short length of the skeletal assembloid at 1 and 6 months post-transplantation. *p < 0.05, **p < 0.01, ***p < 0.001, ****p < 0.0001. (F) µCT image of the skeletal assembloid at 6 months post-transplantation. (G) Engraftment was histologically assessed at 1 month post-transplantation by Safranin O staining and immunohistological analysis. (H) Engraftment was histologically assessed at 6 months post-transplantation by Safranin O staining and immunohistological analysis.

## DISCUSSION

In this study, we successfully developed a strategy to selectively induce two distinct chondrocyte lineage organoids, namely, IZ/ACP and GPC organoids, from hPSCs via ontogenetically-defined ExpLBM cells. Several conventional chondrogenic protocols using BMP2/4 and TGFβ1/3 ^18,30,31^, including ours ^23,24^, have been reported previously and yielded mixtures of both articular chondrocyte and GPC phenotypes (Figure S1B). Here, we discovered that addition of the MEK inhibitor PD0325901 (IZ/ACP organoid strategy) preferentially drives cells toward the “interzone/articular cartilage” lineage capable of forming permanent hyaline cartilaginous tissue without evidence of ossification, whereas elevation of the BMP4 and TGFβ1 concentrations (GPC organoid strategy) promotes generation of the “growth plate cartilage” lineage capable of endochondral ossification, giving rise to bone tissue with vasculature and osteoclasts.

MAPK signaling regulates chondrocyte growth, differentiation, and hypertrophy during endochondral ossification ^32–34^. Although no evidence has previously demonstrated its importance in generating interzone or articular cartilage progenitors, unbiased chemical screening using GDF5-tdTomato reporter hPSCs in this study unveiled this aspect (Figure 1B). In fact, scRNA-seq datasets of the human embryonic limb clearly indicated that MAPK signaling is attenuated during developmental processes from LBM cells to articular cartilage (Figure S3B). Therefore, in conjunction with scRNA-seq data indicating that our organoids exhibit similar signatures as embryonic limb constituent cells (Figures 1H and 1I), we confirmed that this stepwise induction from hPSCs to ExpLBM cells and each differentiated organoid faithfully recapitulates the human developmental program of articular cartilage and growth plate cartilage. Furthermore, in a rat knee defect model, IZ/ACP organoids engrafted into the joint defect area and secreted an extracellular matrix containing ACAN and PRG4, which are characteristics of articular cartilage. Meanwhile, GPC organoids transplanted into a femoral defect model formed cartilage calluses that underwent endochondral ossification, eventually leading to bone union and partial recovery of mechanical strength. Thus, our protocol not only faithfully recapitulates the *in vivo* signaling pathways that drive lineage bifurcation during human limb development but also sheds light on how a promising cell source can be provided for regenerative medicine.

An assembloid is an advanced 3D culture model created by combining or fusing multiple organoids. By integrating different types of organoids or cell populations, assembloids aim to recapitulate more complex tissue and organ-like structures and functions than traditional single-lineage organoids ^35,37^. A further highlight of our study is the generation of a skeletal assembloid by fusing IZ/ACP and GPC organoids. When transplanted into immunodeficient mice, the assembloid exhibited longitudinal growth and endochondral ossification at its center, while maintaining cartilage-like tissues at both ends. This arrangement mimics the limb developmental organization, where growth plate cartilage regions are flanked by articular cartilage regions. While several studies have recapitulated endochondral ossification in cartilage tissue derived from hPSCs ^10–14^, this is the first study in which the process has been faithfully reproduced in a directional manner along the skeletal longitudinal axis. This model holds great promise for investigating complex human skeletal development as well as for studying pathological conditions affecting both cartilage and bone, such as osteoarthritis and skeletal dysplasia.

### Limitations of the study

While this study presents an innovative assembloid approach with hPSC-derived IZ/ACP and GPC organoids to recapitulate human limb skeletal development, it is important to note certain limitations. Longitudinal skeletal growth of the assembloid in this study was observed through subcutaneous transplantation into immunodeficient mice; thus, functional integration of the transplanted tissue within a normal immune environment and under external factors such as mechanical forces has not yet been determined. In fact, we could not observe a secondary ossification center, which typically forms postnatally in the epiphysis of mammals ^36^. Furthermore, given the potential of skeletal assembloids, it is important to utilize this system in future studies to model diseases that cause growth retardation due to impaired limb endochondral ossification.

## RESOURCE AVAILABILITY

### Lead contact

Further information and requests for resources and reagents should be directed to and will be fulfilled by the lead contact, Takeshi Takarada.

### Materials availability

tdTomato reporter hPSCs generated in this study are available from the lead contact with a completed materials transfer agreement.

### Data and code availability

All raw sequencing data were deposited in the GEO database. These data can be accessed using the GSE accession number GSExxxxx (in progress). This study also analyzed publicly available data. Accession numbers of these datasets are listed in the STAR Methods and key resources table. Any additional information and materials will be made available upon request.

## ACKNOWLEDGEMENTS

We thank Ayume Nitta, Kokoro Furutani, Aoba Nakajima, Yukio Hitsumoto, Masayuki Okamoto, Yuka Yamamoto, and Naoshige Ono for technical assistance, and Ziyi Wang and Mitsuaki Ono for their invaluable comments and advice. Preparation of slides and staining were supported by the Central Research Laboratory, Okayama University Medical School. Computation was performed using the General Projects on the supercomputer “Flow” at the Information Technology Center, Nagoya University. We thank the members of the Department of Animal Resources, Advanced Science Research Center, and Okayama University for maintaining mice. This research was supported by Grants-in-Aid for Scientific Research from the Japan Society for the Promotion of Science (17H04399, 21H02643, and 24KK0206 to T.Takarada; and 23K08677 to T.Takao), AMED (JP24bm1123059 to T.Takao), the JST FOREST Program (JPMJFR225H to T.Takarada), the Astellas Foundation for Research on Metabolic Disorders (to T.Takao), and the Cooperative Research Program of the Institute for Frontier Life and Medical Sciences, Kyoto University (to T.Takarada and J.T.). These funders had no role in the study design, data collection and analysis, decision to publish, or preparation of the manuscript.

## AUTHOR CONTRIBUTIONS

T.Takao and T.Takarada conceived the project, designed experiments, and prepared the manuscript. T.O. analyzed sequencing data. K.S. transplanted organoids into critical-sized femur defects in SCID rats. D.Y. and Y.F. performed experiments. M.H. provided critical advice on producing a skeletal assembloid. E.N., T.O., and J.T. discussed the data and provided critical advice.

## STAR METHODS

### KEY RESOURCES TABLE

| REAGENT or RESOURCE | SOURCE | IDENTIFIER |
| --- | --- | --- |
| **Antibodies** |  |  |
| Rabbit polyclonal anti-SOX9 | Sigma-Aldrich | Cat# AB5535<br>RRID: AB_2239761 |
| Rabbit polyclonal anti-ACAN | Proteintech | Cat# 13880-1-AP<br>RRID: AB_2722780 |
| Mouse monoclonal anti-COL1 | SouthernBiotech | Cat# 1441-01<br>RRID: AB_2794758 |
| Mouse monoclonal anti-COL2 | Thermo Fisher Scientific | Cat# MA1-37493<br>RRID: AB_2082337 |
| Mouse monoclonal anti-COL10 | Thermo Fisher Scientific | Cat#14-9771-80<br>RRID: AB_2573017 |
| Rabbit polyclonal anti-PRG4 | Novus Biologicals | Cat# NBP1-19048<br>RRID: AB_2169241 |
| Rabbit monoclonal anti-RUNX2 | CST | Cat# 12556<br>RRID: AB_2732805 |
| Mouse monoclonal anti-SP7 | R&D | Cat# MAB7547<br>RRID: AB_3658831 |
| Rat anti-mouse CD107b (LAMP2) | BD Pharmingen | Cat# 550292<br>RRID: AB_393587 |
| Rabbit monoclonal anti-cathepsin K | Abcam | Cat# ab19027<br>RRID: AB_2261274 |
| Rat monoclonal anti-EMCN | Santa Cruz | Cat# sc-65495<br>RRID: AB_2100037 |
| Rabbit monoclonal anti-hVIM | Abcam | Cat# ab16700<br>RRID: AB_443435 |
| Mouse monoclonal anti-human nucleoli | Abcam | Cat# ab190710<br>RRID: AB_2937055 |
| Rabbit polyclonal anti-KuO | MBL | Cat# PM051M<br>RRID: AB_2876863 |
| Rabbit polyclonal anti-histone H3K27ac | Active Motif | Cat# 39134<br>RRID: AB_2722569 |
| Rabbit polyclonal anti-histone H3K27me3 | Active Motif | Cat# 39155<br>RRID: AB_2561020 |
| Goat anti-mouse IgG (H+L), F(ab')2 fragment (Alexa Fluor 647 conjugate) | CST | Cat# 4410S |
| Goat anti-rabbit IgG (H+L), F(ab')2 fragment (Alexa Fluor 488 conjugate) | CST | Cat# 4412S |
| Goat anti-rabbit IgG (H+L), F(ab')2 fragment (Alexa Fluor 647 conjugate) | CST | Cat# 4414S |
| Goat anti-rat IgG (H+L) (Alexa Fluor 488 conjugate) | CST | Cat# 4416S |
| Goat anti-rat IgG (H+L) (Alexa Fluor 647 conjugate) | CST | Cat# 4418S |
| **Chemicals, peptides, and recombinant proteins** |  |  |
| CHIR99021 | Axon Medchem | Cat# Axon1386 |
| PIK-90 | Selleck | Cat# S1187 |
| A 83-01 | Tocris | Cat# 2939 |
| Wnt-C59 | Cellagen Technology | Cat# C7641-2s |
| LDN-193189 | REPROCELL | Cat# 04-0074 |
| GDC-0449 (Vismodegib) | Cellagen Technology | Cat# C4044-5 |
| Y-27632 | MedChemExpress | Cat# MCE HY-10583 |
| SAG 21k | R&D | Cat# 5282 |
| ATRA | Sigma-Aldrich | Cat# R2625 |
| XAV-939 | Selleck | Cat# S1180 |
| SF1670 | Cayman | Cat# 15368 |
| MK-2206 | Cayman | Cat# 11593 |
| PD0325901 | Selleck | Cat# S1036 |
| VO-OHPic | Cayman | Cat# 10009965 |
| MHY1485 | Cayman | Cat# 18008 |
| DAPT | Wako | Cat# 043-33581 |
| Rapamycin | Selleck | Cat# S1039 |
| Recombinant human activin A | R&D | Cat# 338-AC |
| Recombinant human BMP4 | R&D | Cat# 314-BP |
| Recombinant human TGFβ1 | PEPROTECH | Cat# 100-21C |
| Recombinant human GDF5 | PEPROTECH | Cat# 120-01 |
| Recombinant human FGF2 | Katayama Chemical | Cat# 161-0010 |
| Recombinant human EGF | STEMCELL TECHNOLOGIES | Cat# ST-78006 |
| **Critical commercial assays** |  |  |
| RNeasy Mini Kit | QIAGEN | Cat# 74106 |
| ReverTra Ace qPCR RT Master Mix with gDNA Remover | TOYOBO | Cat# FSQ-301 |
| Luna Universal qPCR Master Mix | NEB | Cat# M3003X |
| KAPA RNA HyperPrep Kit with RiboErase (HMR) | Nippon Genetics | Cat# KK8561 |
| SeqCap Adapter Kit (Set A/B) | Roche | Cat# 07141530001<br>Cat# 07141548001 |
| Chromium Next GEM Single Cell 3' GEM Kit v.3.1 | 10x Genomics | Cat# 1000269 |
| Chromium Next GEM Single Cell G Tip Kit | 10x Genomics | Cat# 1000127 |
| Dual Index Kit TT Set A | 10x Genomics | Cat# 1000215 |
| High sensitivity D5000 screen tape | Agilent | Cat# 5067-5592 |
| Illumina Tagment DNA Enzyme and Buffer | Illumina | Cat# 20034197 |
| MinElute PCR Purification Kit | QIAGEN |  |
| Nextera Index Kit | Illumina | Cat# 15055290 |
| CUT&TAG-IT Assay Kit, Anti-Rabbit | Active Motif | Cat# 53160 |
| RNAscope Multiplex Fluorescent Reagent Kit ver. 2 | Advanced Cell Diagnostics | Cat# 323100 |
| GFP probe (RNAscope) | Advanced Cell Diagnostics | Cat# 538851 |
| TSA Vivid Fluorophore 570 (RNAscope) | Advanced Cell Diagnostics | Cat# 323272 |
| **Deposited data** |  |  |
| Raw and analyzed data (bulk RNA-seq, scRNA-seq, ATAC-seq, and CUT&Tag) | This study | GSExxxxx (in progress) |
| Human limb-bud data (scRNA-seq) | Zhang et al. <sup>27</sup> | E-MTAB-8813 |
| Stepwise differentiation data from hiPSCs into cartilage (bulk RNA-seq) | Yamada et al. <sup>23</sup> | GSE165620 |
| **Experimental models: Cell lines** |  |  |
| 414C2 hiPSCs | CiRA, Kyoto University | N/A |
| SEES4 hESCs | RIKEN BRC | HES0017 |
| 802-3G hiPSCs | REPROCELL | RCRP004N |
| GDF5<sup>3tdTomato</sup> reporter 414C2 hiPSCs | This paper | N/A |
| SP7<sup>3tdTomato</sup> reporter SEES4 hESCs | This paper | N/A |
| CAG-GFP hiPSCs | This paper | N/A |
| CAG-KuO hiPSCs | This paper | N/A |
| **Experimental models: Organisms/strains** |  |  |
| NOD-SCID mouse (NOD.Cg-*Prkdc*<sup>scid</sup>/J) | The Jackson Laboratory, Japan | N/A |
| C57BL/6J mouse | CLEA Japan | N/A |
| SCID rat (F344-*Il2rg*<sup>em11exas</sup>) | Laboratory Animal Research, The University of Tokyo | Cat# 12076-002 |
| **Oligonucleotides** |  |  |
| Primers for RT-qPCR and genotyping, see Table S1 | This paper | N/A |

| Recombinant DNA |  |  |
| --- | --- | --- |
| pPB-CAG-DEST-pA-pgk-bsd | Addgene | Cat# 74918 |
| KW158 (PCAG-PBase) | Kim et al.<sup>38</sup> | Kindly provided by Dr. Knut Woltjen |
| pSpCas9(BB)-2A-Puro (PX459) V2.0 | Addgene | Cat# 62988 |
| PX459-GDF5 gRNA | This paper | N/A |
| PX459-SP7 gRNA | This paper | N/A |
| pUCFa-GDF5 3'HA HA-IRES-tdTomato-PGK-Neo | This paper | N/A |
| pTA2-SP7 3'HA HA-IRES-tdTomato-PGK-Neo | This paper | N/A |
| Software and algorithms |  |  |
| GraphPad Prism 8 | GraphPad software | https://www.graphpad.com/ |
| Fastp (version 0.23.4) | OpenGene | https://github.com/OpenGene/fastp |
| Kallisto (version 0.48.0) | Pachter laboratory | https://pachterlab.github.io/kallisto/ |
| Bowtie2 (version 2.2.5) | Quinlan laboratory | https://bowtie-bio.sourceforge.net/bowtie2/index.shtml |
| Gopeaks (version 1.0.0) | Maxson Braun laboratory | https://github.com/maxsonBraunLab/gopeaks |
| Bedtools (version 2.30.0) | Quinlan laboratory | https://github.com/arq5x/bedtools2 |
| Homer (version 4.11) | Benner laboratory | http://homer.ucsd.edu/homer/ |
| deeptools (version 3.5.1) | Max Planck Institute | https://github.com/deeptools/deepTools |
| Cell Ranger (version 7.0.1) | 10x Genomics | https://www.10xgenomics.com/jp/support/software/cell-ranger/latest |
| R (version 4.1.3) | R Foundation | https://www.r-project.org/ |
| edgeR (version 4.0.3) | Smyth laboratory | https://github.com/OliverVoogd/edgeR |
| NOISeq (version 2.46.0) | Conesa laboratory | https://github.com/ConesaLab/NOISeq |
| Seurat (version 5.0.3) | Satija laboratory | https://satijalab.org/seurat/ |
| DoubletFinder (version 2.0.4) | Git Hub | https://github.com/chris-mcginnis-ucsf/DoubletFinder |
| Other |  |  |
| StemFit | Ajinomoto | Cat# AK02N |
| DMEM | Wako | Cat# 048-29763 |
| IMDM, GlutaMAX_ supplement | Thermo Fisher Scientific | Cat# 31980097 |
| Ham's F-12 Nutrient Mix, GlutaMAX_ supplement | Thermo Fisher Scientific | Cat# 1765092 |
| Polyvinyl alcohol | Sigma-Aldrich | Cat# P8136-250G |
| Chemically defined lipid concentrate (CD lipid concentrate) | Thermo Fisher Scientific | Cat# 11905031 |
| Monothioglycerol | Sigma-Aldrich | Cat# M6145 |
| Insulin | Sigma-Aldrich | Cat# 11376497001 |
| Transferrin | Roche | Cat# 10652202001 |
| L(+)-Ascorbic acid | Wako | Cat# 014-04801 |
| Insulin-transferrin-selenium (ITS) 100x | Thermo Fisher Scientific | Cat# 41400045 |
| Fetal bovine serum (FBS) | Gibco | Cat# 10270106 (Lot: 42F5094K) |
| iMatrix-511 silk (iMatrix) | MATRIXOME | Cat# 892021 |
| Human plasma fibronectin (1 mg/mL) | Sigma-Aldrich | Cat# FC010 |
| Penicillin-streptomycin (10,000 U/mL) | Thermo Fisher Scientific | Cat# 15140122 |
| G418 | Life Technologies | Cat# 076-05962 |
| TrypLE Select | Thermo Fisher Scientific | Cat# 12563029 |
| Accutase | NACALAI TESQUE | Cat# 12679-54 |
| Collagenase, type II, powder | Gibco | Cat# 17101015 |
| 0.5 M EDTA (pH 8.0) solution | NIPPON GENE | Cat# 311-90075 |
| 96-well clear ultra-low attachment microplates | CORNING | Cat# 7007 |
| Agarose, low gelling temperature | Sigma-Aldrich | Cat# A9414-5G |
| 4',6-diamidino-2-phenylindole (DAPI) | Thermo Fisher Scientific | Cat# D3571 |
| Calcein-AM | DOJINDO | Cat# C326 |
| In-Fusion HD Cloning Kit | Takara | Cat# Z9648N |
| KOD FX NEO Kit | Takara | Cat# KFX-201 |
| pTA2 vector (TARget Clone™) | TOYOBO | Cat# TAK-101 |
| 4% paraformaldehyde (PFA) | Wako | Cat# 163-20145 |
| Hematoxylin | Wako | Cat# 131-09665 |
| Eosin | Wako | Cat# 051-06515 |
| Safranin O | Wako | Cat# 530-00592 |
| Fast green | Wako | Cat# 061-00031 |
| Silver nitrate | Wako | Cat# 195-09382 |
| Entellan™ new | Merck | Cat# 1079610100 |
| Normal goat serum (NGS) | Vector Laboratories | Cat# S-1000-20 |
| Fluoromount-G | SouthernBiotech | Cat# 0100-01 |

### EXPERIMENTAL MODEL

#### Animals

The experiments involving mice and rats were approved by the Okayama University Animal Care and Use Committee. Immunodeficient NOD-SCID mice and SCID rats were purchased from The Jackson Laboratory (Japan) and Laboratory Animal Research (The University of Tokyo), respectively. The animals were housed under a 12-h light/12-h dark cycle (06:00–18:00) in a temperature-controlled room maintained at 22 ± 1°C with 40–60% humidity. Food and water were provided ad libitum.

#### Cell lines

##### hPSC lines and maintenance

hPSCs were cultured and maintained with StemFit (Ajinomoto). Before reaching subconfluency, cells were dissociated with TrypLE Select (Thermo Fisher Scientific)/0.25 mM EDTA and suspended in StemFit containing 10 μM Y-27632 (MedChemExpress). Cells (1 × 10^4^) were suspended in StemFit containing 10 µM Y-27632 and 8 µL of iMatrix (human laminin-511 E8 fragment, MATRIXOME) and added to a 6-cm dish. The next day, culture media were replaced with fresh StemFit lacking Y27632. Culture media were changed every 2 days until the next passage. The following hPSC lines were used: 414C2 hiPSCs (provided by the Center for iPS Cell Research and Application, Kyoto University, Japan) and 802-3G hiPSCs (REPROCELL). SEES4 hESCs were donated by RIKEN BRC.

##### Generation of GDF5^3’tdTomato^ and SP7^3’tdTomato^ reporter cell lines

The targeting strategy for knock-in of the IRES-tdTomato-PGK-Neo cassette at the *GDF5* 3′-UTR or *SP7* 3′-UTR or to achieve bicistronic expression of GDF5/SP7 and tdTomato is shown in Figure S2A. The *GDF5* 3′ homology region (GDF5 3′HA) was obtained by an artificial gene synthesis service (pUCFa-GDF5 3′HA vector) (FASMAC). To construct the targeting vector (pUCFa-GDF5 3′HA HA-IRES-tdTomato-PGK-Neo), IRES-tdTomato-PGKNeo fragments were inserted into the pUCFa-GDF5 3′HA vector using an In-Fusion HD Cloning Kit (Takara). Guide RNAs (gRNAs) were designed to target the PAM sequence located 161 bp downstream from the stop codon of GDF5 (CTGAGTGTGACTTGGGCTAA<u>AGG</u>, underlining indicates the PAM sequence). gRNA oligonucleotides (Table S1) were designed, phosphorylated, annealed, and subcloned into the PX459 vector (Addgene) harboring a Cas9 expression cassette (PX459-GDF5 gRNA). The *SP7* 3′ homology region (SP7 3′HA) was synthesized using PCR with a KOD FX NEO Kit (Takara). Genomic DNA extracted from 414C2 hiPSCs, which served as a template, was ligated with the pTA2 vector (TOYOBO) by TA cloning (pTA2-SP7 3′HA vector). To construct the targeting vector (pTA2-SP7 3′HA HA-IRES-tdTomato-PGK-Neo), IRES-tdTomato-PGKNeo fragments were inserted into the pTA2-SP7 3′HA vector using an In-Fusion HD Cloning Kit (Takara). gRNAs were designed to target the PAM sequence located 104 bp downstream from the stop codon of *SP7* (CCCATGCATGCCATCCTTCG<u>GGG</u>, underlining indicates the PAM sequence). gRNA oligonucleotides (Table S1) were designed, phosphorylated, annealed, and subcloned into the PX459 vector (Addgene) harboring a Cas9 expression cassette (PX459-SP7 gRNA).

To generate the GDF5^3′tdTomato^ or SP7^3′tdTomato^ reporter line, 1 × 10^6^ 414C2 hiPSCs (GDF5^3′tdTomato^ reporter) or SEES4 hESCs (SP7^3′tdTomato^ reporter) were electroporated with the targeting vector (7.5 μg) and Cas9/gRNA expression vector (2.5 μg). Selection with G418 (Life Technologies) was maintained for 10 days until stable colonies appeared. Colonies with diameters >500 μm were selected using a P10 pipette tip with the aid of a microscope and used to inoculate larger volumes of media. To verify precise integration of the IRES-tdTomato-PGKNeo cassette into the GDF5/SP7 3′UTR locus, the 5′ or 3′ ends between GDF5/SP7 and the IRES-tdTomato-PGKNeo cassette were amplified for nucleotide sequence analysis. The PCR primers used are listed in Table S1.

##### Generation of CAG-GFP and CAG-KuO reporter cell lines

To generate the CAG-GFP hiPSC and CAG-KuO hiPSC reporter lines, 1 × 10^6^ single cells dissociated from 414C2 hiPSCs were electroporated with 5 μg of the piggyBac Transposase expression plasmid (PCAG-PBase, kindly provided by Dr. Knut Woltjen) ^38^ and 5 μg of the donor plasmid (pPB-CAG-GFP or pPB-CAG-KuO) using Super Electroporator NEPA21 Type II (NEPAGENE), and then plated onto iMatrix-coated plates. Three days after electroporation, cells were treated with puromycin for 1 week and allowed to recover for another 5–7 days. Then, GFP^+^ or KuO^+^ clones were picked and expanded.

### METHOD DETAILS

#### Generation of ExpLBM cells from hPSCs

For stepwise differentiation into LBM cells ^23,24^, hPSCs (3 × 10^4^) were suspended in 1 mL of StemFit (Ajinomoto) containing 10 µM Y-27632 (MedChemExpress), and 4 µL of iMatrix was added to a 3.5-cm culture dish. The next day, the culture medium was replaced with fresh StemFit lacking Y27632. After 2 days of culture, cells were washed with phosphate-buffered saline (PBS) and differentiation was induced by changing to different culture medium (based on chemically defined CDM2 medium) at each time point as described previously ^24^. In brief, cells were treated with mid-primitive streak (MPS) medium (CDM2 basal medium containing 30 ng/mL activin A, 40 ng/mL BMP4, 6 µM CHIR99021, 100 nM PIK-90, and 10 µM Y27632) for 1 day to induce MPS cells (day 1), LPM medium (CDM2 basal medium containing 1 µM A 83-01, 30 ng/mL BMP4, 1 µM C59, and 10 µM Y-27632) for 1 day to induce LPM cells (day 2), and finally LBM medium (CDM2 basal medium containing 1 µM A 83-01, 0.5 µM LDN-193189, 3 µM CHIR99021, 150 nM Vismodegib, and 10 µM Y-27632) for 2 days to induce LBM cells (day 4). The composition of CDM2 basal medium was 50% IMDM (with GlutaMAX, HEPES, and sodium bicarbonate; Gibco) and 50% F12 (with GlutaMAX) containing 1 mg/mL polyvinyl alcohol (Sigma-Aldrich), 1% CD lipid concentrate (Gibco), 450 μM monothioglycerol (Sigma-Aldrich), 7 μg/mL insulin (Sigma-Aldrich), 15 μg/mL transferrin (Roche), and 1% (v/v) penicillin/streptomycin (Thermo Fisher Scientific). To generate ExpLBM cells, LBM cells were dissociated using accutase (NACALAI TESQUE), and 2 × 10^5^ cells were suspended in ExpLBM medium [CDM2 basal medium containing 3 µM CHIR99021, 1 µM A 83-01, and 20 ng/mL FGF2 (Katayama Chemicals); 20 ng/mL EGF (STEMCELL TECHNOLOGIES); and 10 µM Y-27632] and cultured on a 6-cm dish coated with 4 μg/mL human plasma fibronectin (Sigma-Aldrich). The culture media was replaced with fresh ExpLBM medium every 2 days. Before reaching subconfluency, the cells were passaged as described above. The details of the human recombinant proteins and small-molecule agonists and inhibitors are provided in the key resources table.

#### Chondrogenic induction (generation of IZ/ACP organoids, GPC organoids, and skeletal assembloids)

To perform chondrogenic induction of ExpLBM cells under pellet culture conditions (3DCI), 2 × 10 ExpLBM cells were suspended in 200 µL of step 1 medium [CDM2 medium containing 3 µM CHIR99021, 10 ng/mL FGF2, and 50 µg/mL ascorbic acid (Wako) and 1× ITS (Thermo Fisher Scientific)] and plated into 96-well ultra-low U-bottom plates (CORNING). The cells were pelleted by centrifugation at 2,000 rpm for 5 min. After 6 days of culture, the culture medium was changed to step 2 medium [CDM2 medium containing 50 µg/mL ascorbic acid, 1× ITS, and 30 ng/mL BMP4 (R&D); 10 ng/mL TGFβ1 (PEPROTECH); 10 ng/mL GDF5 (PEPROTECH); and 10 ng/mL FGF2] and cultured for an additional 6 days. Subsequently, the medium was replaced with step 3 medium (CDM2 medium containing 50 µg/mL ascorbic acid, 1× ITS, 30 ng/mL BMP4, 10 ng/mL TGFβ1, and 10 ng/mL GDF5) and maintained for 14 days in the presence of various tested chemicals. To generate IZ/ACP organoids, step 3 medium supplemented with 500 nM PD0325901 was used (#16 condition, Figure 1B). GPC organoids were generated using CDM2 medium containing 50 µg/mL ascorbic acid, 1× ITS, 300 ng/mL BMP4, 100 ng/mL TGFβ1, and 10 ng/mL GDF5 during the 14-day step 3 period (#2 condition, Figure 1B). The culture medium was refreshed with fresh medium every 3 days. To generate skeletal assembloids, two IZ/ACP organoids were positioned vertically at both ends of a GPC organoid within low-melting point agarose gel (Sigma-Aldrich) and then cultured in DMEM (Wako) supplemented with 10% FBS and 1% (v/v) penicillin/streptomycin (Thermo Fisher Scientific) for 7 days to allow spontaneous fusion of these samples. The media were changed every other day.

#### Bulk RNA-seq and data processing

Total RNA was extracted using a RNeasy Mini Kit (QIAGEN), and sequencing libraries were prepared using a KAPA RNA HyperPrep Kit with RiboErase (HMR) (Nippon Genetics) and a SeqCap Adapter Kit (Set A or Set B) (Roche) following the manufacturers’ instructions. Sequencing was performed using NovaSeq 6000 (Illumina) with a pair-read sequencing length of 150 bp. All sequence reads were extracted in FASTQ format using the CASAVA 1.8.4 pipeline. Fastp (v0.23.4) was used to remove adapters and filter raw reads with < 60 bases in addition to leading and trailing bases with a read quality of less than 30. For pseudoalignment of the filtered reads to GRCh38, Kallisto (v0.48.0) was used. GeTMM normalization was performed using edgeR (v4.0.3) to account for sample variation. Differentially expressed genes were identified by NOISeq (v2.46.0) analysis with a threshold of p > 0.8 and abs(Log2FC) > 1. For conversion of hiPSCs to ExpLBM cells, our previous data were reanalyzed (GSE165620).

#### ATAC-seq and data processing

ATAC-seq was conducted following a previously described method ^39^. In brief, 4000 cells were collected and lysed using lysis buffer containing 10 mM Tris-HCl, 10 mM NaCl, 3 mM MgCl_2_, and 0.1% IGEPAL CA-630. The Tn5 transposase reaction using Illumina Tagment DNA Enzyme and Buffer (Illumina) was performed at 37°C for 30 min. The reacted DNA was purified using a MinElute PCR Purification Kit (QIAGEN) and amplified for 8–15 cycles using a Nextera Index Kit (Illumina) to produce libraries for sequencing. Sequencing was performed using HiSeq1500 (Illumina) with a pair-read sequencing length of 150 bp. Two or three biological replicates were analyzed. Sequences were aligned to human genome sequences (hg38) using Bowtie2 with default parameters. Gopeaks was used to call peaks using default parameters. Regions where overlapping peaks were identified by Bedtools that intersected between biological replicates were designated as high-confidence peaks. The read count matrix of each sample was used to detect differentially accessible peaks using edgeR. To generate a heatmap, normalized read counts obtained using edgeR were z-score-scaled and plotted. For motif analysis, findMotifsGenome.pl of Homer was used with the -size200-mask option. For annotation of peaks, annotatePeaks.pl of Homer was used with default settings. For visualization, BAM files were converted to bigwig format using deeptools with the command “bamCoverage -binSize 1 - normalizeUsing RPGC --extendReads”.

#### CUT&Tag analysis of histone modifications and data processing

A total of 20,000 cells were collected as input, and libraries were prepared using a CUT&Tag-IT Assay Kit (Active Motif) following the manufacturer’s protocol. Anti-H3K27ac (Active Motif) and anti-H3K27me3 (Active Motif) antibodies were used for reactions. Three biological replicates were analyzed. Sequencing was performed using HiSeq1500 (Illumina) with a pair-read sequencing length of 150 bp. Sequences were aligned to human genome sequences (hg38) using Bowtie2 (parameters: --end-to-end, --very-sensitive, --no-mixed, --no-discordant, -- phred33, -I 10, -X 700). For peak calling, gopeaks and bedtools were utilized as described in the previous section. BAM files were converted to bigwig format using deeptools with the command “bamCoverage -binSize 1 -normalizeUsing RPGC --extendReads”. A heatmap was created using the “computeMatrix” and “plotHeatmap” functions of deeptools. To separate promoter and non-promoter regions, the “annotatePeaks.pl” function of Homer was utilized, and a promoter was defined as one that was annotated as a promoter-TSS in the default setting.

#### *In vivo* transplantation

For subcutaneous transplantation, organoids and assembloids were transplanted into immunodeficient NOD-SCID mice. For transplantation into osteochondral defects created in knee joint articular cartilage, the joint capsules of the knee joints of 10-week-old SCID rats (F344-Il2rgem1Iexas) were opened. A drill hole with a 1-mm diameter and a 0.5-mm depth was created at the femoral groove. Organoids were transplanted into the osteochondral defect of articular cartilage via press fitting. The joint capsules were closed, and rats were sacrificed 3 months later. For transplantation into critical-sized femur defects of SCID rats, a critical-size defect measuring 5 mm was created in the midshaft of the femur using a drill. The created defect was stabilized using an external fixator, and large GPC organoids were implanted into the defect. For postoperative evaluation of bone regeneration, radiographic imaging and histological analysis were performed at 1 and 3 months after surgery.

Transplanted tissues were harvested and fixed with 4% PFA (Wako) for 24 h. Tissue samples were paraffin-embedded at the Central Research Laboratory, Okayama University Medical School. For hematoxylin and eosin (HE) staining, deparaffinized sections were stained with hematoxylin (Wako) for 20 min and with eosin (Wako) for 1 min. For Safranin O staining, deparaffinized sections were stained with 0.05% Fast Green (Wako) solution for 5 min and then with 0.1% Safranin O (Wako) solution for 5 min. For von Kossa staining, fixed samples were incubated in 5% silver nitrate (Wako) under sunlight for 60 min and then treated with 5% sodium thiosulfate for 3 min. After staining, samples were dehydrated and embedded with Entellan new reagent (Merck). Images were acquired using a BZ-X710 microscope (Keyence).

#### Flow cytometry

Following dissociation with accutase, 1 × 10^5^ cells were suspended in 100 µL of PBS containing 2% FBS and incubated with an antibody (diluted 1:200) for 1 h on ice. Fluorescence was detected using a CytoFLEX S flow cytometer (Beckman Coulter), and data were analyzed using FlowJo software (FlowJo LLC). The antibodies used are listed in the key resources table.

#### RT-qPCR

Total RNA was extracted using a RNeasy Mini Kit, and cDNA was synthesized using ReverTra Ace qPCR RT Master Mix with gDNA Remover (TOYOBO) according to the manufacturer’s protocol. cDNAs were then used as templates for qPCR analysis with Luna Universal qPCR Master Mix (NEB) and gene-specific primers. qPCR was performed using an AriaMX Real-Time PCR System (Agilent). The cycle parameters were as follows: denaturation at 95°C for 30 sec, annealing at 62°C for 30 sec, and elongation at 72°C for 30 sec. The expression level of each gene was calculated using the 2^−ΔΔCt^ method. The primer sequences are provided in Table S1.

#### Immunohistochemistry

Tissues were fixed with 4% PFA, and paraffin-embedded samples were sectioned (4 μm thick). Tissue samples were deparaffinized, and antigens were activated by heating slides in 10 mM citrate buffer (pH 6.0). Samples were incubated with blocking solution [1× PBS (−) containing 3% NGS (Vector Laboratories) and 0.1% Triton X-100]. Samples were treated with a primary antibody (diluted 1:200) overnight at 4°C and were incubated with blocking solution [1× PBS (−) containing 3% NGS and 0.1% Triton X-100], followed by a secondary antibody (diluted 1:400) for 1 h at room temperature. Samples were embedded with Fluoromount-G (SouthernBiotech) after staining with DAPI (Thermo Fisher Scientific), and images were acquired using a BZ-X710 camera. The antibodies are listed in the key resources table.

#### scRNA-seq and data processing

ExpLBM cells were washed with PBS (−) and dissociated with accutase at 37°C for 5 min, while IZ/ACP and GPC organoids were washed with PBS (−) and then treated with 900 U/mL collagenase (type II) (Gibco) for 60 min at 37°C, followed by gentle pipetting using a serological pipette to dislodge and dissociate aggregates. Cells were sorted on a FACSAria III Cell Sorter (BD Biosciences) to remove dead cells and debris via forward scatter, side scatter, and DAPI/calcein-AM staining. A total of 10,000 cells were analyzed. A Chromium Next GEM Single Cell 3′ GEM Kit v3.1 (10x Genomics) was used for library preparation. Dissociated cells were separated into single droplets with unique molecular barcodes and then processed following the manufacturer’s protocol. Quality checks of the samples were performed using high sensitivity D5000 screen tape (Agilent) and the Agilent 4150 TapeStation system (Agilent). Sequencing was performed using DNBseq (MGI).

To obtain count matrices, reads were aligned to the human genome GRCh38 using Cell Ranger (v7.0.1) with default parameters. Downstream analysis was performed using the R package Seurat (v5.0.3) ^40^. For quality control, cells with more than five detected genes, a UMI count higher than 5, and <10% mitochondrial genes were kept as high-quality cells. Doublets were detected using DoubletFinder (v2.0.4) ^41^ and removed. Counts were normalized, and then the top 3000 highly variable genes were detected via the SCTransform and FindVariableFeatures functions of Seurat.

For data integration, the batch effect was removed by the LeverageScore method via SketchData in Seurat. After scaling gene expression and performing dimensional reduction, cells were visualized using UMAP embedding. Cluster annotations were labeled with canonical markers.

Transcriptome similarity was compared with published developing human limb [5–9 postconceptional weeks (PCW)] scRNA-seq data using anchor-based label transfer implemented in Seurat. To this end, anchors between the query and reference dataset were identified and filtered using the FindTransferAnchors function with the top 14 dimensions. Then, the prediction score was calculated using the TransferData function. The cell type and region similarity (defined as the mean prediction score of each cell type) were visualized in a heatmap.

Trajectory analysis was performed using the monocle3 package (v1.3.7) ^42^ with default parameters, and the Mes1 cluster was chosen as the root node to order cells. A pseudotime heatmap was plotted using the ComplexHeatmap (v2.18.0) package.

#### SEM and TEM

For SEM, samples were fixed overnight at 4°C in 0.1 M phosphate buffer (pH 7.4) containing 2% PFA and 2% glutaraldehyde. After washing, samples were post-fixed with 2% osmium tetroxide for 90 min at room temperature. Following washing, tissues were dehydrated using a series of ascending ethanol solutions (30%, 50%, 70%, 90%, and two changes of 100%) for 10, 10, 10, 30, 30, and 30 min, respectively. Dehydrated tissues were immersed in t-butyl alcohol for 30 min and freeze-dried using an EIKO ID-2 freeze dryer. Dried samples were cracked with a razor blade to expose the photoreceptor layer and mounted on the aluminum sample stage. Sample surfaces were coated with osmium using HPC-1SW (vacuum device). Subsequently, samples were analyzed using a Hitachi S-4800 scanning electron microscope. For TEM, cartilaginous particles were fixed with 4% PFA and 2% glutaraldehyde, followed by post-fixation with 2% osmium tetroxide. After dehydration, embedding, and polymerization, ultra-thin sections were stained with 2% uranyl acetate and observed using a Hitachi H-7100S transmission electron microscope at an acceleration voltage of 80 kV.

#### *In situ* hybridization

Paraffin sections were stained for GFP using an RNAscope Multiplex Fluorescent Reagent Kit ver. 2 (Advanced Cell Diagnostics). More precisely, sections were pretreated with RNAScope hydrogen peroxidase for 10 min at room temperature, with RNAScope Target Retrieval for 15 min at 98°C, and with protease plus treatment for 30 min at 40°C. Sections were then hybridized with a GFP probe (Advanced Cell Diagnostics) for 2 h at 40°C and amplified through six sequential steps according to the manufacturer’s instructions. The probe signal was visualized with TSA Vivid Fluorophore 570 (Advanced Cell Diagnostics). Then, slides were observed using a BZ-X710 camera.

#### µCT analysis

To analyze critical-sized femoral defects in SCID rats, imaging was performed using a Latheta LCT-200 (Hitachi Aloka Medical) µCT system with a voltage of 50 kV and a current of 200 µA. To analyze skeletal assembloids, scans were performed on ScanXmate-L090HH (Comscantecno) with a voltage of 74 kV and a current of 100 µA. The resolution was 11.906 µm per pixel. 3D images were reconstructed using VXRE, VX3D (3D Industrial Imaging), and TRI/3D-BON-FCS (RATOC System Engineering).

### QUANTIFICATION AND STATISTICAL ANALYSIS

Data were analyzed using Prism 8. All data were acquired by performing biological replicates of two or three independent experiments and are presented as the mean ± standard error of the mean. Statistical significance was determined using a two-tailed Student’s t-test and an ordinary one-way analysis of variance with correction for multiple testing using Tukey’s method.

---

**Figure S1:**
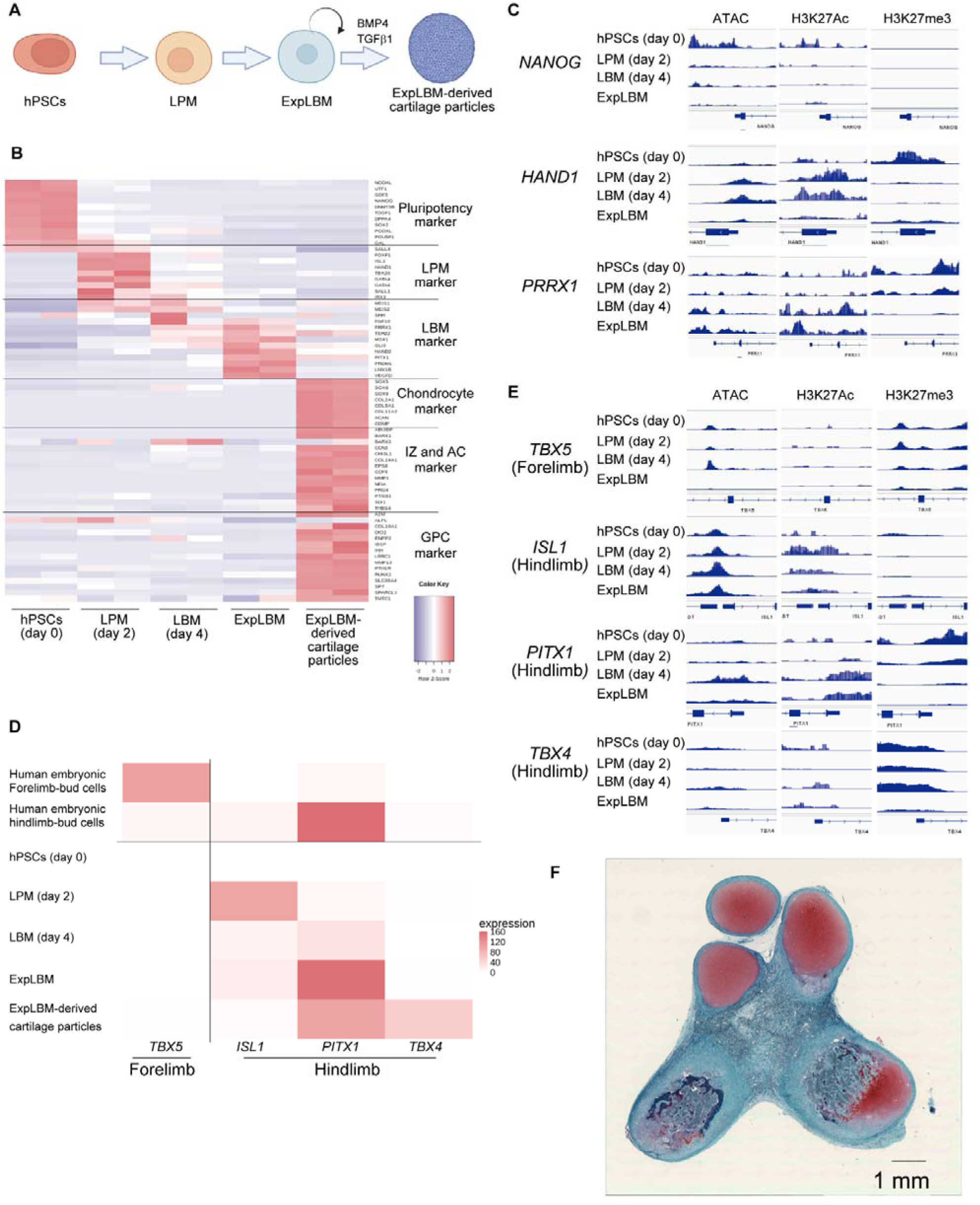
*In vitro* stepwise induction of hPSCs into LBM cells in a manner consistent with hindlimb specification. (A) Schematic view of the hierarchical induction of ExpLBM cells and subsequent differentiation into hyaline cartilaginous-like tissues. (B) Heatmap of gene expression levels of markers in bulk RNA-seq transcriptome analysis of hPSCs (day 0), LPM cells (day 2), LBM cells (day 4), ExpLBM cells (passage numbers 4–5), and ExpLBM-derived cartilage particles generated with 30 ng/mL BMP4, 10 ng/mL TGFβ1, and 10 ng/mL GDF5. n = 2 biologically independent experiments. (C) Integrative Genomics Viewer (IGV) screenshots showing peaks from ATAC-seq and CUT&Tag analyses around marker genes in hPSCs (day 0), LPM cells (day 2), LBM cells (day 4), and ExpLBM cells (passage numbers 4–5). (D) Heatmap showing expression of the forelimb-specific gene *TBX5* and the hindlimb-specific genes *ISL1*, *PITX1*, and *TBX4* in human embryonic forelimb and hindlimb bud cells from scRNA-seq analysis of human embryonic limb cells ^27^, and in bulk RNA-seq transcriptome analysis of hPSCs (day 0), LPM cells (day 2), LBM cells (day 4), ExpLBM cells (passage numbers 4–5), and ExpLBM-derived cartilage particles treated with 30 ng/mL BMP4, 10 ng/mL TGFβ1, and 10 ng/mL GDF5. Data represent the mean of n = 2 biologically independent experiments. (E) IGV screenshots showing peaks from ATAC-seq and CUT&Tag analyses of forelimb and hindlimb marker genes in hPSCs (day 0), LPM cells (day 2), LBM cells (day 4), and ExpLBM cells (passage numbers 4–5). (F) Subcutaneous transplantation of ExpLBM-derived cartilage particles treated with 30 ng/mL BMP4, 10 ng/mL TGFβ1, and 10 ng/mL GDF5 into NOD-SCID mice. Their engraftment was histologically assessed at 5 months post-transplantation by Safranin O staining.

**Figure S2:**
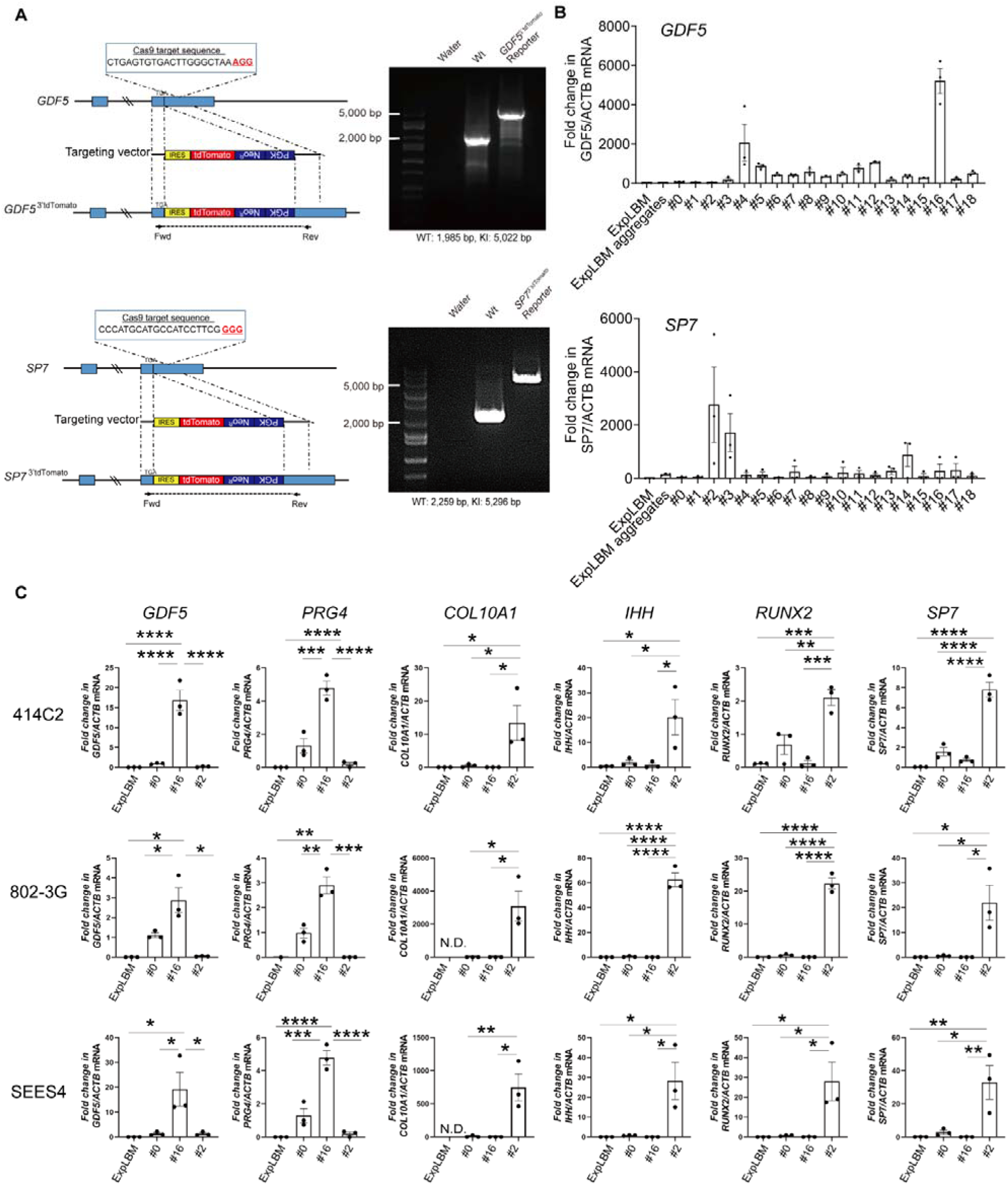
Generation of ontogenetically-defined IZ/ACP and GPC organoids from hPSCs, related to Figure 1. (A) Targeting cassettes of the GDF5^3’tdTomato^ and SP7^3’tdTomato^ knock-in alleles. PAM sequences are highlighted in red. Agarose gel of genotyping PCR of genomic DNA extracted from parental hPSCs and each reporter hPSC line. Targeting cassette sequences were detected using forward and reverse primers that recognize sequences outside the homology arms. WT, wild-type allele; KI, knock-in allele. (B) RT-qPCR analysis of each marker gene in GDF5^3’tdTomato^ and SP7^3’tdTomato^ reporter-derived ExpLBM cells treated with 18 combinations of chemicals and cytokines for 14 days in the presence of 30 ng/mL BMP4 and 10 ng/mL TGFβ1. #0, none; #1, 10% FBS; #2, 300 ng/mL BMP4, 100 ng/mL TGFβ1, and 10% FBS; #3, CHIR99021 (GSK3β inhibitor and WNT activator); #4, C59 (PORCN inhibitor and WNT inhibitor); #5, XAV-939 (tankyrase 1 inhibitor and WNT inhibitor); #6, LDN-193189 (ALK2/3 inhibitor); #7, A 83-01 (ALK4/5/7 inhibitor); #8, SAG 21k (hedgehog signaling activator); #9, Vismodegib (hedgehog signaling inhibitor); #10, MHY1485 (mTOR activator); #11, VO-OHpic (PTEN inhibitor and PI3K/mTOR activator); #12, SF1670 (PTEN inhibitor and PI3K/mTOR activator); #13, rapamycin (mTOR inhibitor); #14, ATRA (RA signaling activator); #15, MK-206 (AKT inhibitor); #16, PD0325901 (MEK inhibitor); #17, PIK-90 (PI3K inhibitor); and #18, DAPT (γ-secretase inhibitor and Notch inhibitor). (C) RT-qPCR analysis of each marker gene in various clones of hPSCs, including 414C2 hiPSCs, 8023-G1 hiPSCs, and SEES4 hESCs. *p < 0.05, **p < 0.01, ***p < 0.001, ****p < 0.0001.

**Figure S3:**
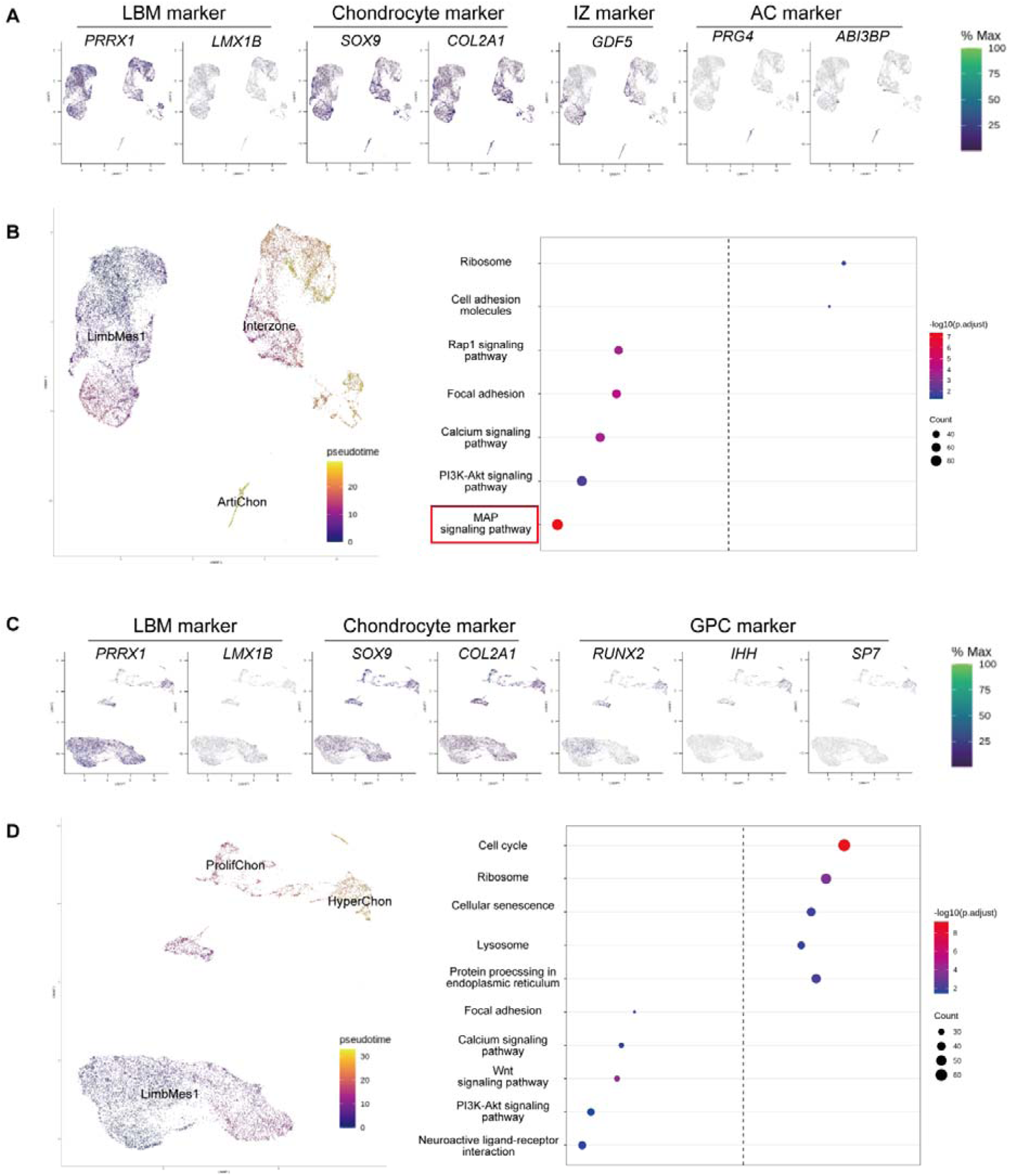
scRNA-seq analysis of human embryonic limb cells, related to Figure 1. (A) Feature plot of each marker gene in the UMAP representation of LimbMes1, Interzone, and ArtiChon clusters, obtained from scRNA-seq datasets of the human embryonic limb. (B) Pseudotime trajectories of subpopulations in the UMAP representation of LimbMes1, Interzone, and ArtiChon clusters, and KEGG pathway enrichment analysis of these subpopulations. (C) Feature plot of each marker gene in the UMAP representation of LimbMes1, ProlifChon, and HyperChon clusters, obtained from scRNA-seq datasets of the human embryonic limb. (D) Pseudotime trajectories of subpopulations in the UMAP representation of LimbMes1, ProlifChon, and HyperChon clusters, and KEGG pathway enrichment analysis of these subpopulations.

**Table S1:**
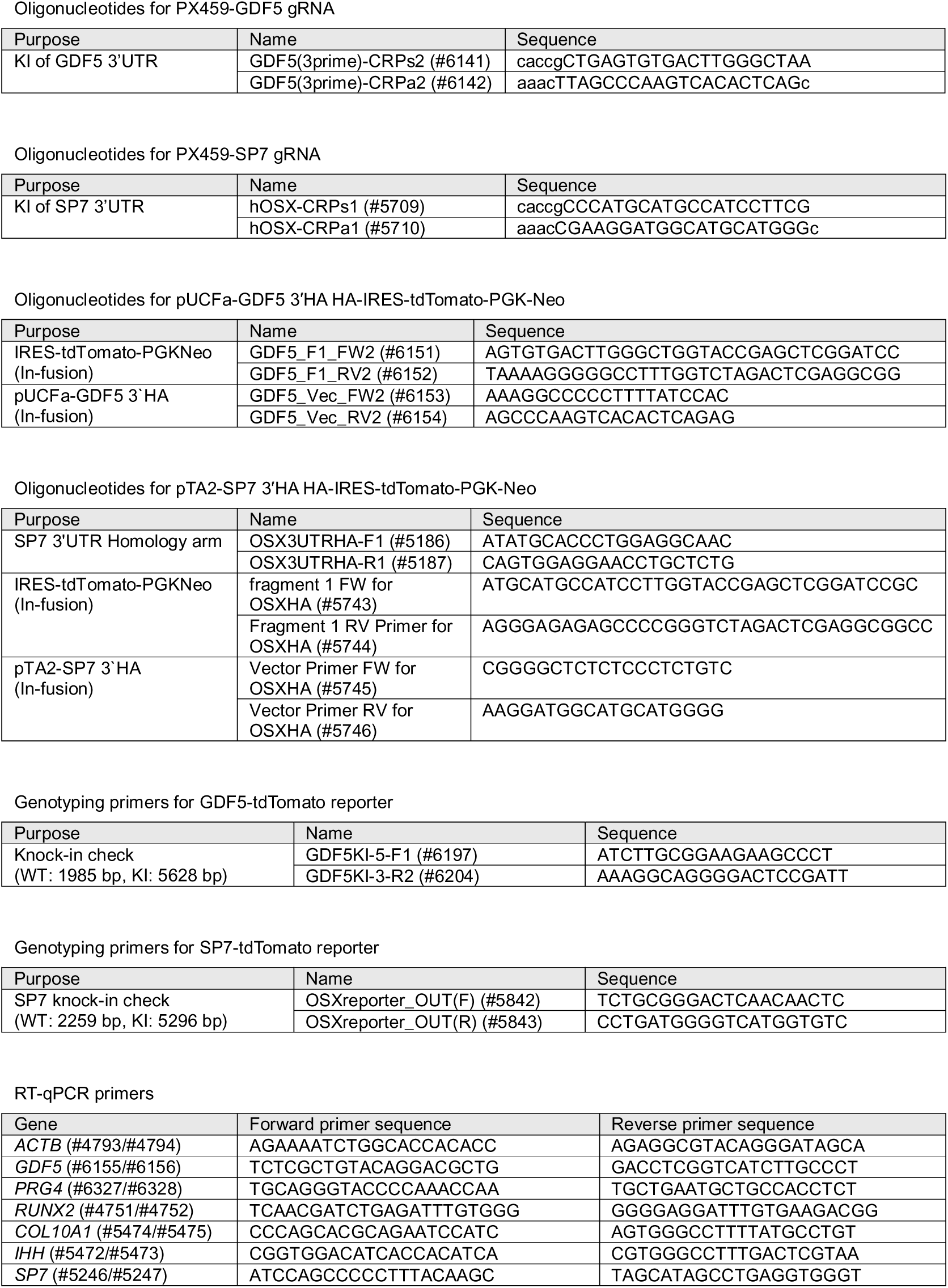
Oligonucleotides used in this study, related to STAR Methods.

## REFERENCES

1. Teti, A. (2011). Bone development: overview of bone cells and signaling. Curr. Osteoporos. Rep. 9, 264–273.

2. Katz, J.N., Arant, K.R., and Loeser, R.F. (2021). Diagnosis and treatment of hip and knee osteoarthritis: A Review: A review. JAMA 325, 568–578.

3. Sharma, L. (2021). Osteoarthritis of the knee. N. Engl. J. Med. 384, 51–59.

4. Calori, G.M., Mazza, E.L., Mazzola, S., Colombo, A., Giardina, F., Romanò, F., and Colombo, M. (2017). Non-unions. Clin. Cases Miner. Bone Metab. 14, 186–188.

5. Geister, K.A., and Camper, S.A. (2015). Advances in skeletal dysplasia genetics. Annu. Rev. Genomics Hum. Genet. 16, 199–227.

6. Yamanaka, S. (2020). Pluripotent stem cell-based cell therapy-promise and challenges. Cell Stem Cell 27, 523–531.

7. Berendsen, A.D., and Olsen, B.R. (2015). Bone development. Bone 80, 14–18.

8. Long, F. (2011). Building strong bones: molecular regulation of the osteoblast lineage. Nat. Rev. Mol. Cell Biol. 13, 27–38.

9. Kronenberg, H.M. (2003). Developmental regulation of the growth plate. Nature 423, 332–336.

10. Ikeda, Y., Tani, S., Moriishi, T., Kuroda, A., Matsuo, Y., Saeki, N., Inui-Yamamoto, C., Abe, M., Maeda, T., Rowe, D.W., et al. (2023). Modeling of intramembranous ossification using human pluripotent stem cell-derived paraxial mesoderm derivatives. Regen Ther 24, 536–546.

11. Kawai, S., Yoshitomi, H., Sunaga, J., Alev, C., Nagata, S., Nishio, M., Hada, M., Koyama, Y., Uemura, M., Sekiguchi, K., et al. (2019). In vitro bone-like nodules generated from patient-derived iPSCs recapitulate pathological bone phenotypes. Nat Biomed Eng 3, 558–570.

12. Kidwai, F., Mui, B.W.H., Arora, D., Iqbal, K., Hockaday, M., de Castro Diaz, L.F., Cherman, N., Martin, D., Myneni, V.D., Ahmad, M., et al. (2020). Lineage-specific differentiation of osteogenic progenitors from pluripotent stem cells reveals the FGF1-RUNX2 association in neural crest-derived osteoprogenitors: FGF1 IN NEURAL CREST-DERIVED OSTEOPROGENITORS. Stem Cells 38, 1107–1123.

13. Xi, H., Fujiwara, W., Gonzalez, K., Jan, M., Liebscher, S., Van Handel, B., Schenke-Layland, K., and Pyle, A.D. (2017). In vivo human somitogenesis guides somite development from hPSCs. Cell Rep. 18, 1573–1585.

14. Kawai, S., Sunaga, J., Nagata, S., Nishio, M., Fukuda, M., Kamakura, T., Sun, L., Jin, Y., Sakamoto, S., Watanabe, A., et al. (2023). 3D osteogenic differentiation of human iPSCs reveals the role of TGFβ signal in the transition from progenitors to osteoblasts and osteoblasts to osteocytes. Sci. Rep. 13, 1094.

15. Loh, K.M., Chen, A., Koh, P.W., Deng, T.Z., Sinha, R., Tsai, J.M., Barkal, A.A., Shen, K.Y., Jain, R., Morganti, R.M., et al. (2016). Mapping the Pairwise Choices Leading from Pluripotency to Human Bone, Heart, and Other Mesoderm Cell Types. Cell 166, 451–467.

16. Wu, C.-L., Dicks, A., Steward, N., Tang, R., Katz, D.B., Choi, Y.-R., and Guilak, F. (2021). Single cell transcriptomic analysis of human pluripotent stem cell chondrogenesis. Nat. Commun. 12, 362.

17. Umeda, K., Zhao, J., Simmons, P., Stanley, E., Elefanty, A., and Nakayama, N. (2012). Human chondrogenic paraxial mesoderm, directed specification and prospective isolation from pluripotent stem cells. Sci. Rep. 2, 455.

18. Craft, A.M., Rockel, J.S., Nartiss, Y., Kandel, R.A., Alman, B.A., and Keller, G.M. (2015). Generation of articular chondrocytes from human pluripotent stem cells. Nat. Biotechnol. 33, 638–645.

19. Pretemer, Y., Kawai, S., Nagata, S., Nishio, M., Watanabe, M., Tamaki, S., Alev, C., Yamanaka, Y., Xue, J.-Y., Wang, Z., et al. (2021). Differentiation of Hypertrophic Chondrocytes from Human iPSCs for the In Vitro Modeling of Chondrodysplasias. Stem Cell Reports 16, 610–625.

20. Tani, S., Okada, H., Onodera, S., Chijimatsu, R., Seki, M., Suzuki, Y., Xin, X., Rowe, D.W., Saito, T., Tanaka, S., et al. (2023). Stem cell-based modeling and single-cell multiomics reveal gene-regulatory mechanisms underlying human skeletal development. Cell Rep., 112276.

21. Iimori, Y., Morioka, M., Koyamatsu, S., and Tsumaki, N. (2021). Implantation of human-induced pluripotent stem cell-derived cartilage in bone defects of mice. Tissue Eng. Part A 27, 1355–1367.

22. Smeeton, J., Askary, A., and Crump, J.G. (2017). Building and maintaining joints by exquisite local control of cell fate. Wiley Interdiscip. Rev. Dev. Biol. 6. 10.1002/wdev.245.

23. Yamada, D., Nakamura, M., Takao, T., Takihira, S., Yoshida, A., Kawai, S., Miura, A., Ming, L., Yoshitomi, H., Gozu, M., et al. (2021). Induction and expansion of human PRRX1+ limb-bud-like mesenchymal cells from pluripotent stem cells. Nat Biomed Eng 5, 926–940.

24. Takao, T., Yamada, D., and Takarada, T. (2022). A protocol to induce expandable limb-bud mesenchymal cells from human pluripotent stem cells. STAR Protoc 3, 101786.

25. Long, H.K., Prescott, S.L., and Wysocka, J. (2016). Ever-changing landscapes: Transcriptional enhancers in development and evolution. Cell 167, 1170–1187.

26. Klemm, S.L., Shipony, Z., and Greenleaf, W.J. (2019). Chromatin accessibility and the regulatory epigenome. Nat. Rev. Genet. 20, 207–220.

27. Zhang, B., He, P., Lawrence, J.E.G., Wang, S., Tuck, E., Williams, B.A., Roberts, K., Kleshchevnikov, V., Mamanova, L., Bolt, L., et al. (2023). A human embryonic limb cell atlas resolved in space and time. Nature. 10.1038/s41586-023-06806-x.

28. McQueen, C., and Towers, M. (2020). Establishing the pattern of the vertebrate limb. Development 147. 10.1242/dev.177956.

29. Baker, C.E., Moore-Lotridge, S.N., Hysong, A.A., Posey, S.L., Robinette, J.P., Blum, D.M., Benvenuti, M.A., Cole, H.A., Egawa, S., Okawa, A., et al. (2018). Bone fracture acute phase response-A unifying theory of fracture repair: Clinical and scientific implications. Clin. Rev. Bone Miner. Metab. 16, 142–158.

30. Yamashita, A., Morioka, M., Yahara, Y., Okada, M., Kobayashi, T., Kuriyama, S., Matsuda, S., and Tsumaki, N. (2015). Generation of scaffoldless hyaline cartilaginous tissue from human iPSCs. Stem Cell Reports 4, 404–418.

31. Umeda, K., Oda, H., Yan, Q., Matthias, N., Zhao, J., Davis, B.R., and Nakayama, N. (2015). Long-term expandable SOX9+ chondrogenic ectomesenchymal cells from human pluripotent stem cells. Stem Cell Reports 4, 712–726.

32. Chen, H., Tan, X.-N., Hu, S., Liu, R.-Q., Peng, L.-H., Li, Y.-M., and Wu, P. (2021). Molecular mechanisms of chondrocyte proliferation and differentiation. Front. Cell Dev. Biol. 9, 664168.

33. Zhang, Y., Pizzute, T., and Pei, M. (2014). A review of crosstalk between MAPK and Wnt signals and its impact on cartilage regeneration. Cell Tissue Res. 358, 633–649.

34. Stanton, L.-A., Underhill, T.M., and Beier, F. (2003). MAP kinases in chondrocyte differentiation. Dev. Biol. 263, 165–175.

35. Onesto, M.M., Kim, J.-I., and Pasca, S.P. (2024). Assembloid models of cell-cell interaction to study tissue and disease biology. Cell Stem Cell 0. 10.1016/j.stem.2024.09.017.

36. Aghajanian, P., and Mohan, S. (2018). The art of building bone: emerging role of chondrocyte-to-osteoblast transdifferentiation in endochondral ossification. Bone Res. 6, 19.

37. Andersen, J., Revah, O., Miura, Y., Thom, N., Amin, N.D., Kelley, K.W., Singh, M., Chen, X., Thete, M.V., Walczak, E.M., et al. (2020). Generation of functional human 3D Cortico-motor assembloids. Cell 183, 1913–1929.e26.

38. Kim, S.-I., Oceguera-Yanez, F., Sakurai, C., Nakagawa, M., Yamanaka, S., and Woltjen, K. (2016). Inducible transgene expression in human iPS cells using versatile All-in-One piggyBac transposons. Methods Mol. Biol. 1357, 111–131.

39. Buenrostro, J.D., Giresi, P.G., Zaba, L.C., Chang, H.Y., and Greenleaf, W.J. (2013). Transposition of native chromatin for fast and sensitive epigenomic profiling of open chromatin, DNA-binding proteins and nucleosome position. Nat. Methods 10, 1213–1218.

40. Hao, Y., Hao, S., Andersen-Nissen, E., Mauck, W.M., 3rd, Zheng, S., Butler, A., Lee, M.J., Wilk, A.J., Darby, C., Zager, M., et al. (2021). Integrated analysis of multimodal single-cell data. Cell 184, 3573–3587.e29.

41. McGinnis, C.S., Murrow, L.M., and Gartner, Z.J. (2019). DoubletFinder: Doublet detection in single-cell RNA sequencing data using artificial nearest neighbors. Cell Syst. 8, 329–337.e4.

42. Trapnell, C., Cacchiarelli, D., Grimsby, J., Pokharel, P., Li, S., Morse, M., Lennon, N.J., Livak, K.J., Mikkelsen, T.S., and Rinn, J.L. (2014). The dynamics and regulators of cell fate decisions are revealed by pseudotemporal ordering of single cells. Nat. Biotechnol. 32, 381–386.

