## Supplementary material for "Modeling human limb skeletal development using human pluripotent stem cell-derived skeletal assembloids": Fig S1

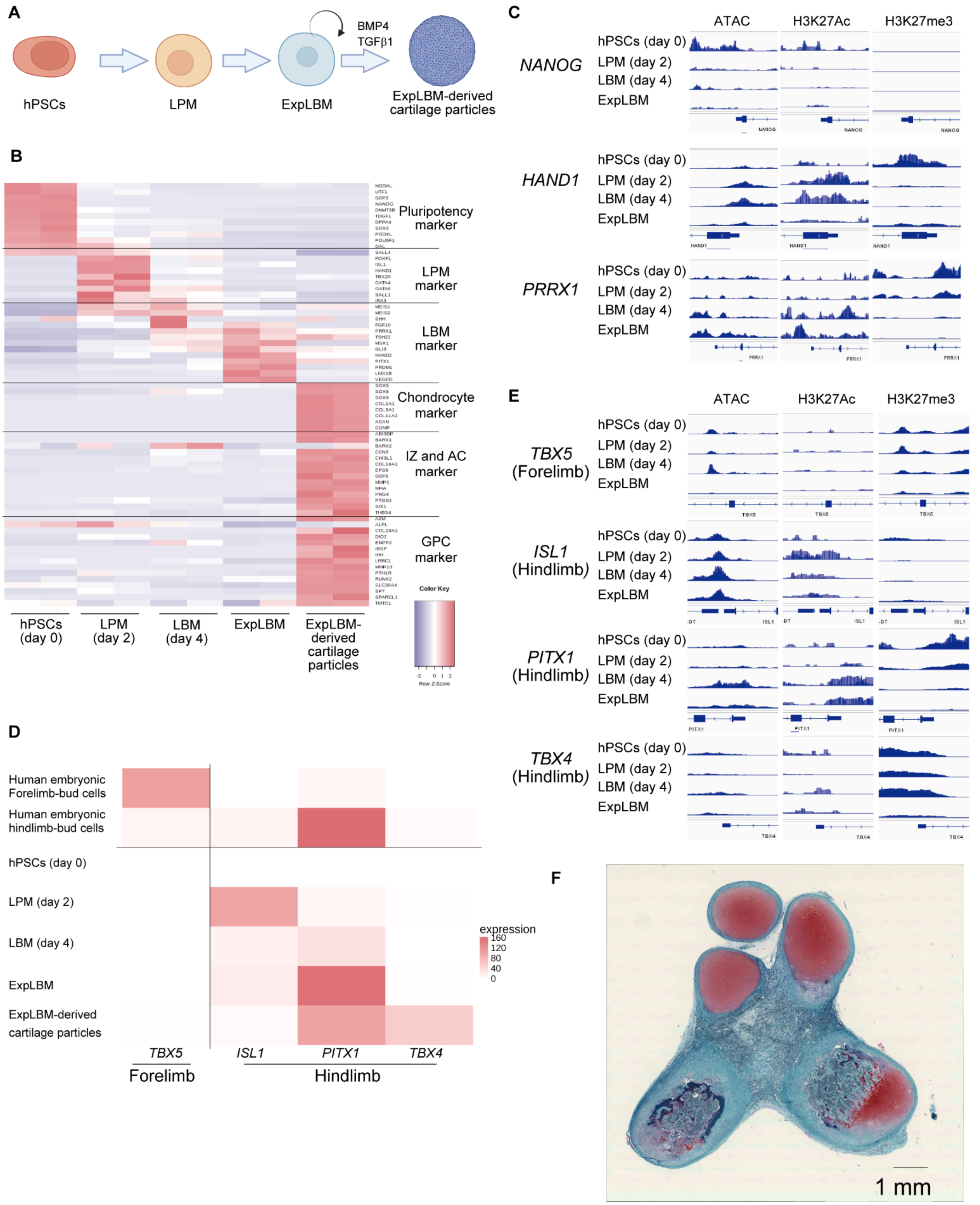
A
hPSCs
LPM
ExpLBM
BMP4
TGFβ1
ExpLBM-derived cartilage particles
B
Pluripotency marker
LPM marker
LBM marker
Chondrocyte marker
IZ and AC marker
GPC marker
Color Key
Row Z-Score
hPSCs (day 0)
LPM (day 2)
LBM (day 4)
ExpLBM
ExpLBM-derived cartilage particles
C
ATAC
H3K27Ac
H3K27me3
NANOG
HAND1
PRRX1
hPSCs (day 0)
LPM (day 2)
LBM (day 4)
ExpLBM
D
Human embryonic Forelimb-bud cells
Human embryonic hindlimb-bud cells
hPSCs (day 0)
LPM (day 2)
LBM (day 4)
ExpLBM
ExpLBM-derived cartilage particles
expression
160
120
80
40
0
TBX5
ISL1
PITX1
TBX4
Forelimb
Hindlimb
E
ATAC
H3K27Ac
H3K27me3
TBX5 (Forelimb)
ISL1 (Hindlimb)
PITX1 (Hindlimb)
TBX4 (Hindlimb)
hPSCs (day 0)
LPM (day 2)
LBM (day 4)
ExpLBM
F
1 mm

Figure S1

**Figure S1: *In vitro* stepwise induction of hPSCs into LBM cells in a manner consistent with hindlimb specification.**

(A) Schematic view of the hierarchical induction of ExpLBM cells and subsequent differentiation into hyaline cartilaginous-like tissues.

(B) Heatmap of gene expression levels of markers in bulk RNA-seq transcriptome analysis of hPSCs (day 0), LPM cells (day 2), LBM cells (day 4), ExpLBM cells (passage numbers 4–5), and ExpLBM-derived cartilage particles generated with 30 ng/mL BMP4, 10 ng/mL TGFβ1, and 10 ng/mL GDF5. n = 2 biologically independent experiments.

(C) Integrative Genomics Viewer (IGV) screenshots showing peaks from ATAC-seq and CUT&Tag analyses around marker genes in hPSCs (day 0), LPM cells (day 2), LBM cells (day 4), and ExpLBM cells (passage numbers 4–5).

(D) Heatmap showing expression of the forelimb-specific gene *TBX5* and the hindlimb-specific genes *ISL1*, *PITX1*, and *TBX4* in human embryonic forelimb and hindlimb bud cells from scRNA-seq analysis of human embryonic limb cells [27], and in bulk RNA-seq transcriptome analysis of hPSCs (day 0), LPM cells (day 2), LBM cells (day 4), ExpLBM cells (passage numbers 4–5), and ExpLBM-derived cartilage particles treated with 30 ng/mL BMP4, 10 ng/mL TGFβ1, and 10 ng/mL GDF5. Data represent the mean of n = 2 biologically independent experiments.

(E) IGV screenshots showing peaks from ATAC-seq and CUT&Tag analyses of forelimb and hindlimb marker genes in hPSCs (day 0), LPM cells (day 2), LBM cells (day 4), and ExpLBM cells (passage numbers 4–5).

(F) Subcutaneous transplantation of ExpLBM-derived cartilage particles treated with 30 ng/mL BMP4, 10 ng/mL TGFβ1, and 10 ng/mL GDF5 into NOD-SCID mice. Their engraftment was histologically assessed at 5 months post-transplantation by Safranin O staining.
