## Supplementary material for "Modeling human limb skeletal development using human pluripotent stem cell-derived skeletal assembloids": Fig S2

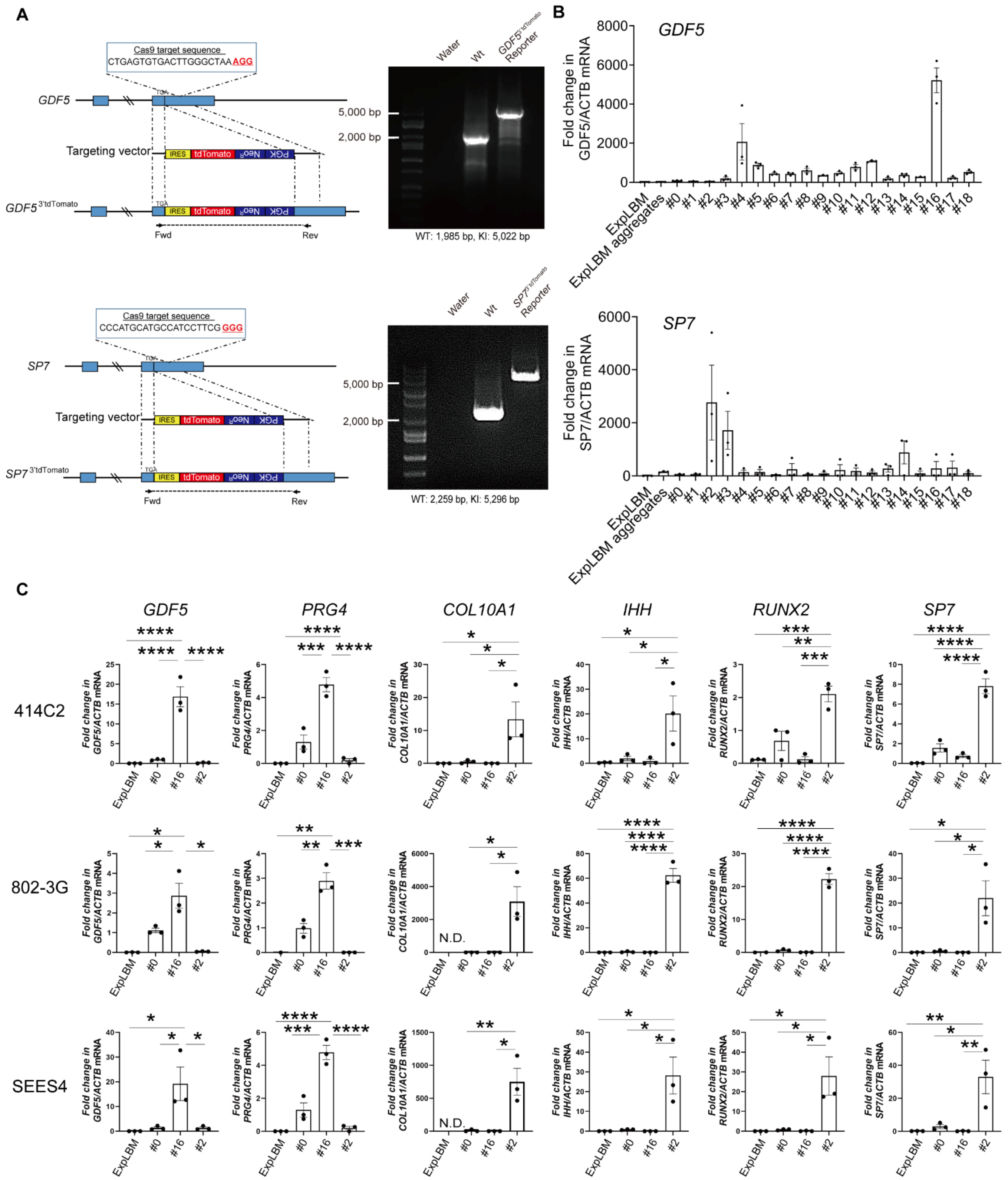
A
Cas9 target sequence
CTGAGTGTGACTTGGGCTAA AGG
GDF5
Targeting vector
IRES tdTomato NeoR PGK
Fwd
Rev
Water
Wt
Reporter
5,000 bp
2,000 bp
WT: 1,985 bp, KI: 5,022 bp
Cas9 target sequence
CCCATGCATGCCATCCTTCG GGG
SP7
WT: 2,259 bp, KI: 5,296 bp
B
GDF5
Fold change in GDF5/ACTB mRNA
SP7
Fold change in SP7/ACTB mRNA
ExpLBM
ExpLBM aggregates
C
414C2
802-3G
SEES4
GDF5
PRG4
COL10A1
IHH
RUNX2
N.D.

Figure S2

**Figure S2: Generation of ontogenetically-defined IZ/ACP and GPC organoids from hPSCs, related to Figure 1.**

(A) Targeting cassettes of the $GDF5^{3'tdTomato}$ and $SP7^{3'tdTomato}$ knock-in alleles. PAM sequences are highlighted in red. Agarose gel of genotyping PCR of genomic DNA extracted from parental hPSCs and each reporter hPSC line. Targeting cassette sequences were detected using forward and reverse primers that recognize sequences outside the homology arms. WT, wild-type allele; KI, knock-in allele.

(B) RT-qPCR analysis of each marker gene in $GDF5^{3'tdTomato}$ and $SP7^{3'tdTomato}$ reporter-derived ExpLBM cells treated with 18 combinations of chemicals and cytokines for 14 days in the presence of 30 ng/mL BMP4 and 10 ng/mL TGFβ1. #0, none; #1, 10% FBS; #2, 300 ng/mL BMP4, 100 ng/mL TGFβ1, and 10% FBS; #3, CHIR99021 (GSK3β inhibitor and WNT activator); #4, C59 (PORCN inhibitor and WNT inhibitor); #5, XAV-939 (tankyrase 1 inhibitor and WNT inhibitor); #6, LDN-193189 (ALK2/3 inhibitor); #7, A 83-01 (ALK4/5/7 inhibitor); #8, SAG 21k (hedgehog signaling activator); #9, Vismodegib (hedgehog signaling inhibitor); #10, MHY1485 (mTOR activator); #11, VO-OHpic (PTEN inhibitor and PI3K/mTOR activator); #12, SF1670 (PTEN inhibitor and PI3K/mTOR activator); #13, rapamycin (mTOR inhibitor); #14, ATRA (RA signaling activator); #15, MK-206 (AKT inhibitor); #16, PD0325901 (MEK inhibitor); #17, PIK-90 (PI3K inhibitor); and #18, DAPT (γ-secretase inhibitor and Notch inhibitor).

(C) RT-qPCR analysis of each marker gene in various clones of hPSCs, including 414C2 hiPSCs, 8023-G1 hiPSCs, and SEES4 hESCs. $*p < 0.05$, $**p < 0.01$, $***p < 0.001$, $****p < 0.0001$.
