## Supplementary material for "Modeling human limb skeletal development using human pluripotent stem cell-derived skeletal assembloids": Fig S3

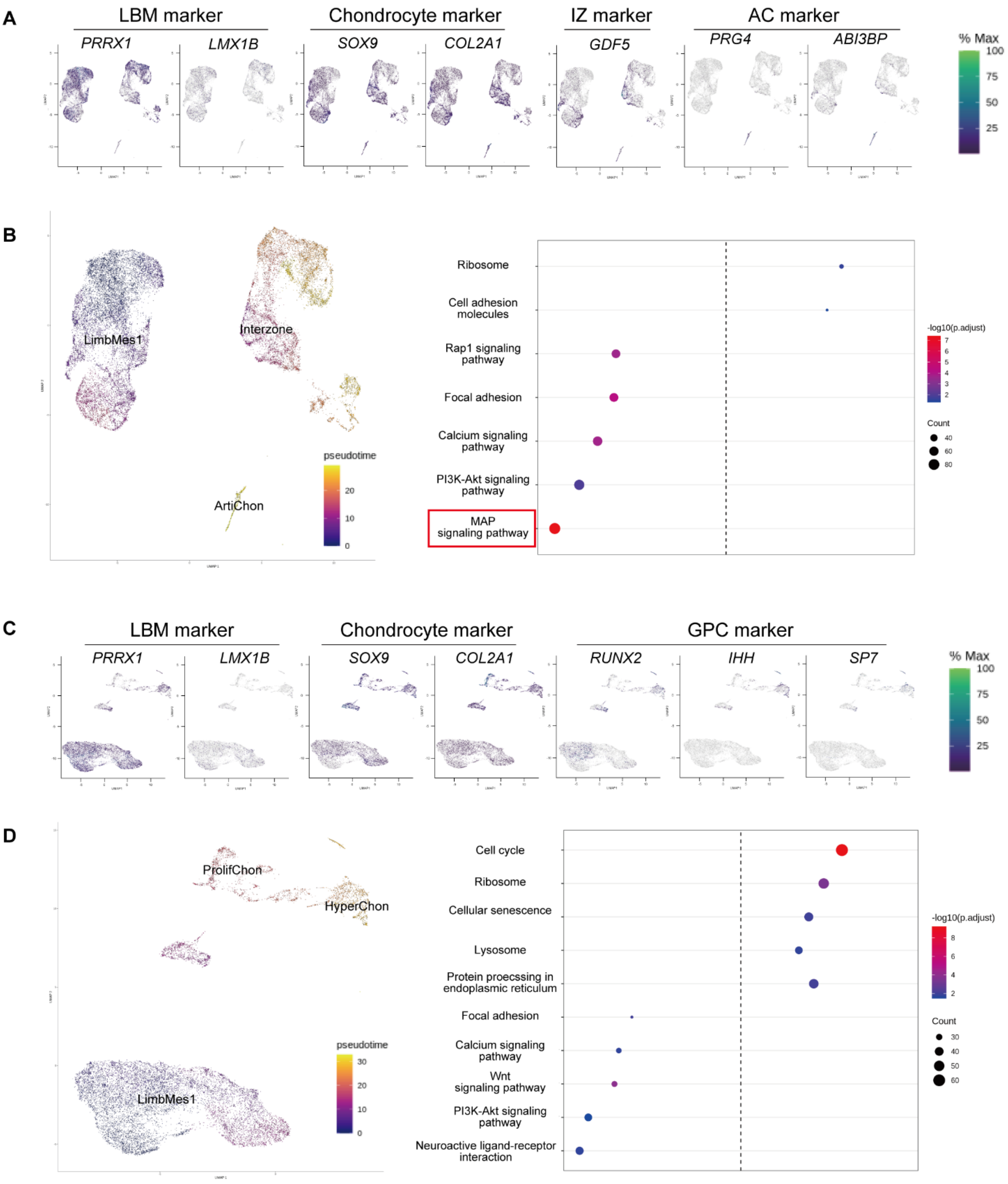
A
LBM marker
Chondrocyte marker
IZ marker
AC marker
PRRX1
LMX1B
SOX9
COL2A1
GDF5
PRG4
ABI3BP
% Max
100
75
50
25
B
LimbMes1
Interzone
ArtiChon
pseudotime
20
10
0
Ribosome
Cell adhesion molecules
Rap1 signaling pathway
Focal adhesion
Calcium signaling pathway
PI3K-Akt signaling pathway
MAP signaling pathway
-log10(p.adjust)
7
6
5
4
3
2
Count
40
60
80
C
LBM marker
Chondrocyte marker
GPC marker
PRRX1
LMX1B
SOX9
COL2A1
RUNX2
IHH
SP7
% Max
100
75
50
25
D
ProlifChon
HyperChon
LimbMes1
pseudotime
30
20
10
0
Cell cycle
Ribosome
Cellular senescence
Lysosome
Protein proecssing in endoplasmic reticulum
Focal adhesion
Calcium signaling pathway
Wnt signaling pathway
PI3K-Akt signaling pathway
Neuroactive ligand-receptor interaction
-log10(p.adjust)
8
6
4
2
Count
30
40
50
60

Figure S3

**Figure S3: scRNA-seq analysis of human embryonic limb cells, related to Figure 1.**

(A) Feature plot of each marker gene in the UMAP representation of LimbMes1, Interzone, and ArtiChon clusters, obtained from scRNA-seq datasets of the human embryonic limb.

(B) Pseudotime trajectories of subpopulations in the UMAP representation of LimbMes1, Interzone, and ArtiChon clusters, and KEGG pathway enrichment analysis of these subpopulations.

(C) Feature plot of each marker gene in the UMAP representation of LimbMes1, ProlifChon, and HyperChon clusters, obtained from scRNA-seq datasets of the human embryonic limb.

(D) Pseudotime trajectories of subpopulations in the UMAP representation of LimbMes1, ProlifChon, and HyperChon clusters, and KEGG pathway enrichment analysis of these subpopulations.
