## Supplementary material for "Modeling human limb skeletal development using human pluripotent stem cell-derived skeletal assembloids": Table S1

### Table S1: Oligonucleotides used in this study, related to STAR Methods.

Oligonucleotides for PX459-GDF5 gRNA

| Purpose | Name | Sequence |
|---|---|---|
| KI of GDF5 3'UTR | GDF5(3prime)-CRPs2 (#6141) | caccgCTGAGTGTGACTTGGGCTAA |
| | GDF5(3prime)-CRPa2 (#6142) | aaacTTAGCCCAAGTCACACTCAGc |

Oligonucleotides for PX459-SP7 gRNA

| Purpose | Name | Sequence |
|---|---|---|
| KI of SP7 3'UTR | hOSX-CRPs1 (#5709) | caccgCCCATGCATGCCATCCTTCG |
| | hOSX-CRPa1 (#5710) | aaacCGAAGGATGGCATGCATGGGc |

Oligonucleotides for pUCFa-GDF5 3′HA HA-IRES-tdTomato-PGK-Neo

| Purpose | Name | Sequence |
|---|---|---|
| IRES-tdTomato-PGKNeo (In-fusion) | GDF5_F1_FW2 (#6151) | AGTGTGACTTGGGCTGGTACCGAGCTCGGATCC |
| | GDF5_F1_RV2 (#6152) | TAAAAGGGGGCCTTTGGTCTAGACTCGAGGCGG |
| pUCFa-GDF5 3`HA (In-fusion) | GDF5_Vec_FW2 (#6153) | AAAGGCCCCCTTTTATCCAC |
| | GDF5_Vec_RV2 (#6154) | AGCCCAAGTCACACTCAGAG |

Oligonucleotides for pTA2-SP7 3′HA HA-IRES-tdTomato-PGK-Neo

| Purpose | Name | Sequence |
|---|---|---|
| SP7 3'UTR Homology arm | OSX3UTRHA-F1 (#5186) | ATATGCACCCTGGAGGCAAC |
| | OSX3UTRHA-R1 (#5187) | CAGTGGAGGAACCTGCTCTG |
| IRES-tdTomato-PGKNeo (In-fusion) | fragment 1 FW for OSXHA (#5743) | ATGCATGCCATCCTTGGTACCGAGCTCGGATCCGC |
| | Fragment 1 RV Primer for OSXHA (#5744) | AGGGAGAGAGCCCCGGGTCTAGACTCGAGGCGGCC |
| pTA2-SP7 3`HA (In-fusion) | Vector Primer FW for OSXHA (#5745) | CGGGGCTCTCTCCCTCTGTC |
| | Vector Primer RV for OSXHA (#5746) | AAGGATGGCATGCATGGGG |

Genotyping primers for GDF5-tdTomato reporter

| Purpose | Name | Sequence |
|---|---|---|
| Knock-in check (WT: 1985 bp, KI: 5628 bp) | GDF5KI-5-F1 (#6197) | ATCTTGCGGAAGAAGCCCT |
| | GDF5KI-3-R2 (#6204) | AAAGGCAGGGGACTCCGATT |

Genotyping primers for SP7-tdTomato reporter

| Purpose | Name | Sequence |
|---|---|---|
| SP7 knock-in check (WT: 2259 bp, KI: 5296 bp) | OSXreporter_OUT(F) (#5842) | TCTGCGGGACTCAACAACTC |
| | OSXreporter_OUT(R) (#5843) | CCTGATGGGGTCATGGTGTC |

RT-qPCR primers

| Gene | Forward primer sequence | Reverse primer sequence |
|---|---|---|
| *ACTB* (#4793/#4794) | AGAAAATCTGGCACCACACC | AGAGGCGTACAGGGATAGCA |
| *GDF5* (#6155/#6156) | TCTCGCTGTACAGGACGCTG | GACCTCGGTCATCTTGCCCT |
| *PRG4* (#6327/#6328) | TGCAGGGTACCCCAAACCAA | TGCTGAATGCTGCCACCTCT |
| *RUNX2* (#4751/#4752) | TCAACGATCTGAGATTTGTGGG | GGGGAGGATTTGTGAAGACGG |
| *COL10A1* (#5474/#5475) | CCCAGCACGCAGAATCCATC | AGTGGGCCTTTTATGCCTGT |
| *IHH* (#5472/#5473) | CGGTGGACATCACCACATCA | CGTGGGCCTTTGACTCGTAA |
| *SP7* (#5246/#5247) | ATCCAGCCCCCTTTACAAGC | TAGCATAGCCTGAGGTGGGT |
